# Somatostatine interneurons mediate dopamine-dependent sensory gating in the orbitofrontal cortex

**DOI:** 10.1101/2025.03.14.643299

**Authors:** Anushree Tripathi, Tilda Sköld, Luciano Censoni, Olof Lagerlöf, Paolo Medini

**Author notes:** Corresponding Authors*: P.Medini and A. Tripathi.

## Abstract

The prefrontal cortex continuously filters irrelevant stimuli, and deficits in this sensory gating predispose to psychotic disorders. In schizophrenia, the normally observed suppression of second response to two paired sounds is drastically reduced. Apart from sensitivity to critical schizophrenia players as NMDA and dopamine, the microcircuitry behind gating remains unknown. We found that the first sound excitatory response recruits, via NMDA receptors, local inhibition to gate the second response via GABAB receptors. Optogenetic inhibition of somatostatine-interneurons counteracted gating more effectively compared to parvalbumine-positive interneurons. Dopamine local blockade also counteracted gating, and dopamine effect was prevented by somatostatine-, but not parvalbumine-, optoactivation. Thus, prefrontal gating is mediated by NMDA-dependent recruitment of dendrite-targeting, somatostatine-interneurons, facilitated by dopamine. The results identify a critical synapse to better target psychose-prone disorders.

## INTRODUCTION

The prefrontal cortex is central for executive functions, controlling lower-level cognitive strategies to accomplish daily goals. A prerequisite for effective executive control is that goal-irrelevant sensory inputs are filtered out by sensory gating (SG). SG is the expression of the inhibitory component of attention, and indeed recent works in mice highlighted the role of different populations of inhibitory interneurons in attentional mechanisms (e.g. (*1*), while clinical research showed links between attentional and SG deficits on the other side(*2*). Psychiatric disorders prone to psychoses like schizophrenia and bipolar disorder are characterized by altered processing of externally as well internally generated neuronal inputs (percepts and thoughts, respectively) leading to hallucinations and delusions, respectively. Recent works highlighted common genetic risk between bipolar disorder and schizophrenia(*3*), an association that is stronger when psychotic bipolar patients are isolated from non-psychotic ones.(*4*). It is nevertheless not easy to find easily measurable physiological readouts of such biological risk, and of the basic deficits in input processing, with one notable exception. Indeed, one of the most reliable, worldwide-recognized endophenotype for biological risk of schizophrenia is SG deficit. SG is a simple electroencephalographic measurement of the local field potential (LFP) response to repeatedly presented sound pairs. In normal subjects, the second sound response is reduced, whereas this “gating” is severely reduced in schizophrenic patients(*5*), and their first-line relatives(*6*), having a higher risk of schizophrenia. SG deficit is more prevalent in psychotic bipolar patients compared to the non-psychotics (*7*), suggesting that SG deficit is an indicator of biological vulnerability to psychosis. On the other hand, altered sensory processing can also be mediating some negative and cognitive symptoms, particularly in the attentional domain, observed in psychotic disorders(*8*) (but see(*9*)). Association between SG and prefrontal executive function has also been documented in normal controls (*10*). Thus, a better understanding of the microcircuit mediating SG can provide insights on which synaptic and molecular mechanisms render prefrontal circuits prone to input processing disturbances contributing to psychosis and executive dysfunction.

Mice are suitable for microcircuit studies, and SG in experimental rodents has been reported in the cortex(*11*) and hippocampus(*12*). Rodent models of schizophrenia also consistently show impaired sensory-motor gating (“pre-pulse inhibition”), like patients, and many drugs able to induce or correct schizophrenia-like symptoms modulate SG accordingly (*13*). For example, NMDA antagonists counteract SG and can cause schizophrenia-like behaviour, and manipulation of catecholaminergic transmission also affects SG, as shown by amphetamine effects (*14*), although the effects of dopaminergic blockade are somewhat controversial(*12*). Nicotinic modulators also affect SG (although the literature is controversial(*12*)), and nicotinic receptors modulate prefrontal GABAergic transmission(*15*), involved in SG via GABAB receptors(*16*). However, the synaptic and microcircuit mechanisms mediating SG remain elusive. To address this question, we used the mouse orbitofrontal cortex (OFC) as a model circuit, as it displays strong local SG(*17*). Our results indicate that SG is mediated by local GABAergic innervation, mainly provided by dendrite-targeting, somatostatin (SS)-positive GABAergic interneurons, and that SS-interneurons also mediate to a significant extent the effect of dopaminergic modulation on SG.

## RESULTS

### Local inhibitory gating of acoustic responses in mouse orbitofrontal cortex

Acoustic responses in ventro-orbital OFC are driven by the acoustic association cortex (AAC)(*17*) When we performed an acoustic SG protocol(*18*), we found that acoustic responses are dramatically gated in OFC but not in its input area AAC (Figure S1A). Figure 1A shows examples and grand-averages (left, middle plots) of LFP response during SG protocol in AAC and OFC (N=6). The amplitudes of 1st and 2nd responses were statistically undistinguishable in AAC (paired t-test, p=0,635), whereas the 2nd response was dramatically smaller in OFC (paired t-test, p=0,004). The statistical comparison of SG ratios confirmed a dramatic SG in OFC compared to AAC (Figure S1B; paired t-test, p=0,001).

**Figure 1.**
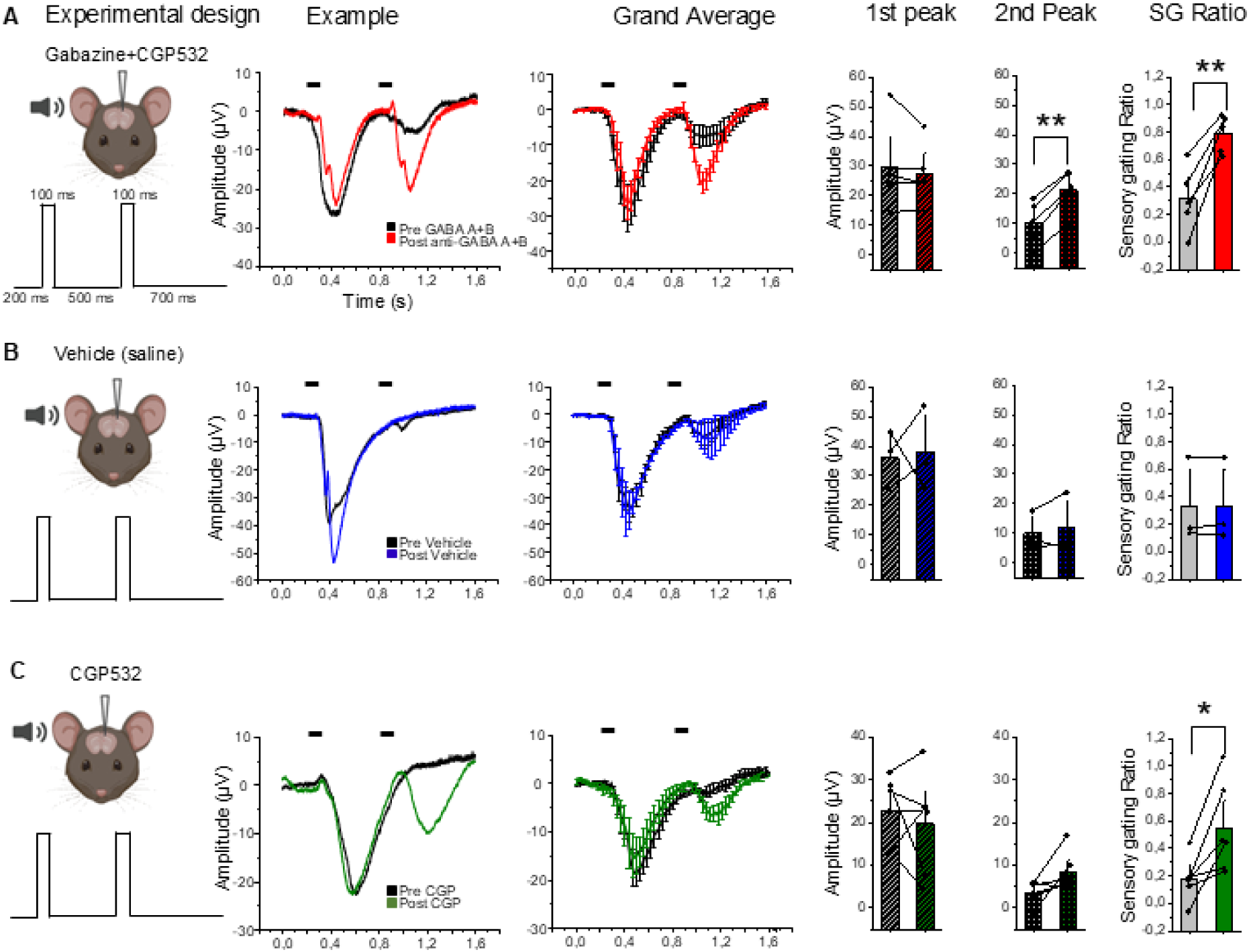
Acoustic SG is driven by local GABAergic circuits in mouse OFC: role of GABA_B_ receptors. **A.** Local infusion of a subepileptic cocktail of GABA_A_ and GABA_B_ antagonists gabazine and CGP 52432 (1,5 and 1µM, respectively) reduced SG by reinstating responsiveness to the second sound without affecting the first response (1^st^ response-pre drug: 29,7 ± 6,6μV *vs* post drug: 27,3 ± 4,6μV; paired t-test, p=0,329; 2^nd^ response-pre drug: 10,1 ± 3,4μV *vs* post drug: 21,3 ± 3,1μV; paired t-test, **p<0,001), resulting in increased SG ratio (pre drug: 0,31± 0,11 *vs* post drug: 0,79 ± 0,06; paired t-test, **p=0,003). From left to right: experiment sketch, examples of pre- and post-injection SG (black and blue, respectively), corresponding grand averages, quantification of 1^st^ and 2^nd^ peak responses and SG ratio. **B.** Same measurements as in **A** for control injections with physiological solution (vehicle of drugs). No change in all measured parameters were detected (1^st^ response-pre drug: 36,05 ± 5,7μV *vs* post drug: 38,05 ± 8,2μV; paired t-test, p=0,868; 2^nd^ response-pre drug: 10,11 ± 3,8μV *vs* post drug: 11,82 ± 6,0μV; paired t-test, p=0,572). No statistical change in the SG ratios-pre vehicle: 0,33 ±0,18 *vs* post vehicle: 0,33 ± 0,18; paired t-test, p=0,839). **C.** Same measurements as in **A,B** upon injection of the sole GABA_B_ antagonist (CGP 52432), which by itself increases SG (from left to right: 1^st^ response-pre drug: 22,7 ± 3,3μV *vs* post drug: 19,8 ± 4,8μV; paired t-test, p=0,53; 2nd response-pre drug: 3,5 ± 1,1μV vs post drug: 8,3±1,9μV; paired t-test p = 0,08; SG ratio-pre drug: 0,18 ± 0,06 *vs* post drug: 0,55 ± 0,13; paired t-test, *p=0,015).

We observed a longer response duration in OFC compared to AAC (Figure S1C; t-test, p=0,013).

We tested whether SG was present in other association areas as the posterior parietal cortex, that also projects to OFC(*19*): no SG was measured as 1^st^ and 2^nd^ peaks were undistinguishable (Figure S1D, N=3; paired t-test, p=0,733). Noticeably, SG was clearly detectable in the frontal association area (FrA in Figure S1E, N=5, paired t-test, p=0,025), indicating that the microcircuit behind SG is prefrontal-specific.

As AAC is the main input area mediating acoustic responses in OFC(*17*), we reasoned that gating of the 2nd response can hardly be attributable to reduced excitatory input from AAC. We tested instead whether SG happens via local GABAergic interneurons within OFC. To test this possibility, we compared SG before and 30 min after a local, intracortical injection of a subepileptic dose of GABA_A_ and GABA_B_ antagonists through the same recording pipette kept in place and at slightly negative pressure pre-injection (Figure 1A-C). These recordings showed a clear increase of the 2nd response compared to pre-injection values, surprisingly without a significant increase of the 1st response (Figure 1A; N=5, 1^st^ peak: paired t-test, p=0,329; 2^nd^ peak: paired t-test, p<0,001).GABA antagonism was thus able to counteract SG as GABA antagonism increased SG ratios(Figure 1A paired t-test, p=0,003).

Vehicle injection, however, failed to show any effect, not even at trend level (Figure 1B; N=3; 1^st^ peak: paired t-test, p=0,868; 2^nd^ peak: paired t-test, p=0,572; SG ratio: paired t-test, p=0,839). As previous work showed that GABA_B_ receptors can modulate hippocampal SG(*16*),we tested whether injection of the sole GABA_B_ antagonist CGP52432 (1μM) was effective in counteracting SG. Figure 1C shows that GABA_B_ antagonism significantly increased the SG ratio (N=6; 1^st^ peak: paired t-test, p=0,53; 2nd peak: paired t-test p=0,08; SG ratio: paired t-test, p=0,015). SG ratios upon GABA_A_ and GABA_B_ antagonism were not different from the SG ratios upon antagonism of the sole GABA_B_ receptor (t-test: p=0,16), indicating that GABA_B_ receptors significantly contributed to gate the 2^nd^ response.

Thus,SG in mouse OFC is largely due to recruitment of a local inhibitory mechanism with a significant contribution from GABA_B_ receptors.

### The first acoustic excitatory response of pyramidal cells drives inhibitory gating of the second one

We next wanted to test whether the first excitatory response drives activation of the inhibitory mechanism that in turn gates the second one. To test this hypothesis we expressed the powerful inhibitory opsin GTCAR1(*20*) in excitatory neurons, by co-injecting a mix of two AAVs, one with a floxed, soma-targeted opsin, and the second coding for the CRE recombinase under the CamKII promotor for excitatory pyramidal neurons. This allowed us to optogenetically reduce excitatory neuron activation selectively during the first pulse (200-800ms; Figure 2A). The examples and grand-averages of Figure 2B show an opsin-driven reduction of first response during opto-inhibition, accompanied by an increase of 2^nd^ response (Figure 2C for data set, N=7; 1^st^ peak: paired t-test, p=0,02; 2^nd^ peak: paired t-test, p=0,03), causing a significant increase of the SG ratio (Figure 2C, paired t-test, p=0,008). We controlled that the 2^nd^ peak increase was not simply due to a rebound excitation of pyramidal neurons due to sudden switch-off of their photoinhibition. Switching off the optic fiber (without any sound presentation) was not able *per se* to induce a detectable downward LFP signal (Figure S2).

**Figure 2.**
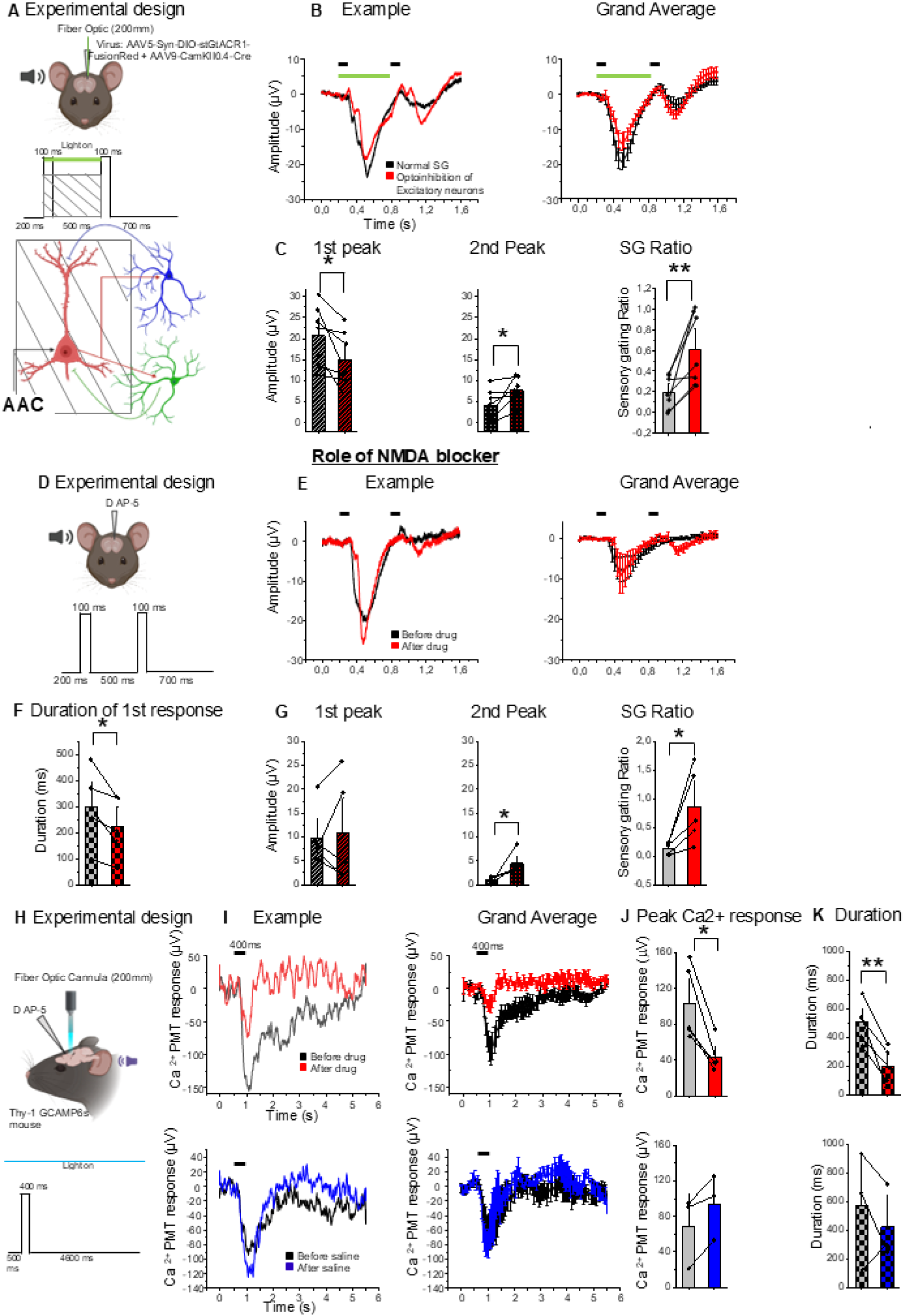
Reduction of the first excitatory response, including its NMDA calcium component, disrupts SG. **A.** Experiment sketch: excitatory pyramidal neurons were optogenetically inhibited during the 1^st^ sound time window while SG was measured electrophysiologically. **B.** Examples and grand averages of LFP responses with the fiber off and on (black and red, respectively). Note the reduction of the 1^st^ response and the increase of the 2^nd^ one. **C.** From left to right: paired quantifications of the 1^st^, 2^nd^ response and of the SG ratio with optogenetic stimulation on and off (black and red, respectively; left – 1^st^ response: 20,7 ± 2,7 µV *vs* after optoinhibition: 14,9 ± 2,5µV; paired t-test, *p=0,02, middle – 2^nd^ response: 4,2 ± 1,4µV *vs* after optoinhibition: 7,7 ± 1,1µV; paired t-test, *p=0,03; right-SG ratios: 0,19 ± 0,06 *vs* after optoinhibition: 0,61± 0,13; paired t-test, **p=0,008). **D.** Experiment sketch. SG was measured before and after local infusion of the NMDA antagonist D-AP5 (0,6μl, 2.5mM) through the recording pipette. **E.** Black and red curves in the examples (left) and gran-averages (right) are measured before and after the drug injection. **F.** Sound response duration, measured as duration at half peak, became shorter upon NMDA antagonism (before NMDA blocker: 299,4 ± 63,2ms *vs* after NMDA blocker: 221,4 ± 52,2ms; paired t-test, *p=0,02). **G.** Quantification of the 1^st^ response (before NMDA blocker: 9,4 ± 2,3 µV *vs* after NMDA blocker: 10,2 ± 4,0µV; paired t-test, p=0,76), 2^nd^ response (before NMDA blocker: 1,1 ± 0,2 µV *vs* after NMDA blocker: 4,0 ± 0,9µV; paired t-test, p=0,05) and of the SG ratio, which was increased by the pharmacological manipulation (before NMDA blocker: 0,14 ± 0,04 *vs* after NMDA blocker: 0,76 ± 0,26; paired t-test, *p=0,04). **H.** Experiment sketch of the calcium fiberphotometric experiment: a blue light emitting optic fiber (0,5NA; 200μm diameter, 0,9mW power, 460nm wavelength) collected the green emission light from Thy1-GCAMP6s transgenic mice. A pipette on a side craniotomy was injecting the NMDA antagonist AP-5. **I.** Examples and grand averages of calcium-related photomultiplier voltage signals recorded in response to acoustic stimulation (single pulse= before and after injection of AP-5 (red lines, top row) or of vehicle (blue lines, saline, bottom row): note the peak reduction upon AP-5 injection and the slight but not significant increase upon saline injection. Examples and grand averages from left to right. **J**. Paired quantifications of the sound-evoked calcium response showed a decrease in response amplitude after injection of NMDA antagonist but not after vehicle injection (before AP-5: 102,2 ± 18,5µV*vs* after AP-5: 43,3 ± 8,0µV; paired t-test, *p=0,01; after saline a non-significant trend for increase was observed: before saline-68,8 ± 13,8µV *vs* after saline-93,5 ± 21,3µV; paired t-test, p=0,06). **K**. The duration of the sound-evoked calcium response (expressed as duration at half peak amplitude) was also significantly reduced after injection of NMDA antagonist but not after vehicle injection (before drug: 502,1 ± 63,3ms *vs* after drug: 196,6 ± 57,1ms; paired t-test, **p=0,004; before saline 570,0 ± 238,1ms *vs* after saline 423,3 ± 149,0ms; paired t-test, p=0,486).

Thus, selective photo-inhibition of pyramidal cells during the 1^st^ response increases the amplitude of the 2^nd^ response, thereby counteracting SG. These results, together with those of the GABA antagonism series, indicate that the excitation of pyramidal neurons in response to 1^st^ sound is what recruits the local inhibitory mechanism that gates the 2^nd^ response.

### NMDA antagonism reduces acoustic response duration and dendritic calcium transients while counteracting SG

Antagonizing glutamate NMDA receptors can induce dissociative and negative symptoms similar to schizophrenia, as well as impair SG(*21*), contributing to the hypothesis of reduced NMDA receptor function in schizophrenia. We thus tested whether NMDA antagonism counteracted SG also in mouse OFC. Injection through the recording pipette of a NMDA selective antagonist D-AP5 puff (Figure 2D; 600nl, 2,5mM(*22*)) counteracted SG (Figure 3A). The examples and grand-averages of Figure 2E show that 2^nd^response increased upon NMDA antagonism, in absence of clear reduction of the first response (paired comparisons pre- and after injections in Figure 2G, left and middle; N=5; 1^st^ paired t-test, p=0,76; 2^nd^ peak: paired t-test, p=0,05). As a result, the SG ratio became significantly higher upon NMDA antagonism (Figure 2G, right, paired t-test, p=0,04), indicating that NMDA antagonism counteracts SG in mouse OFC.

**Figure 3.**
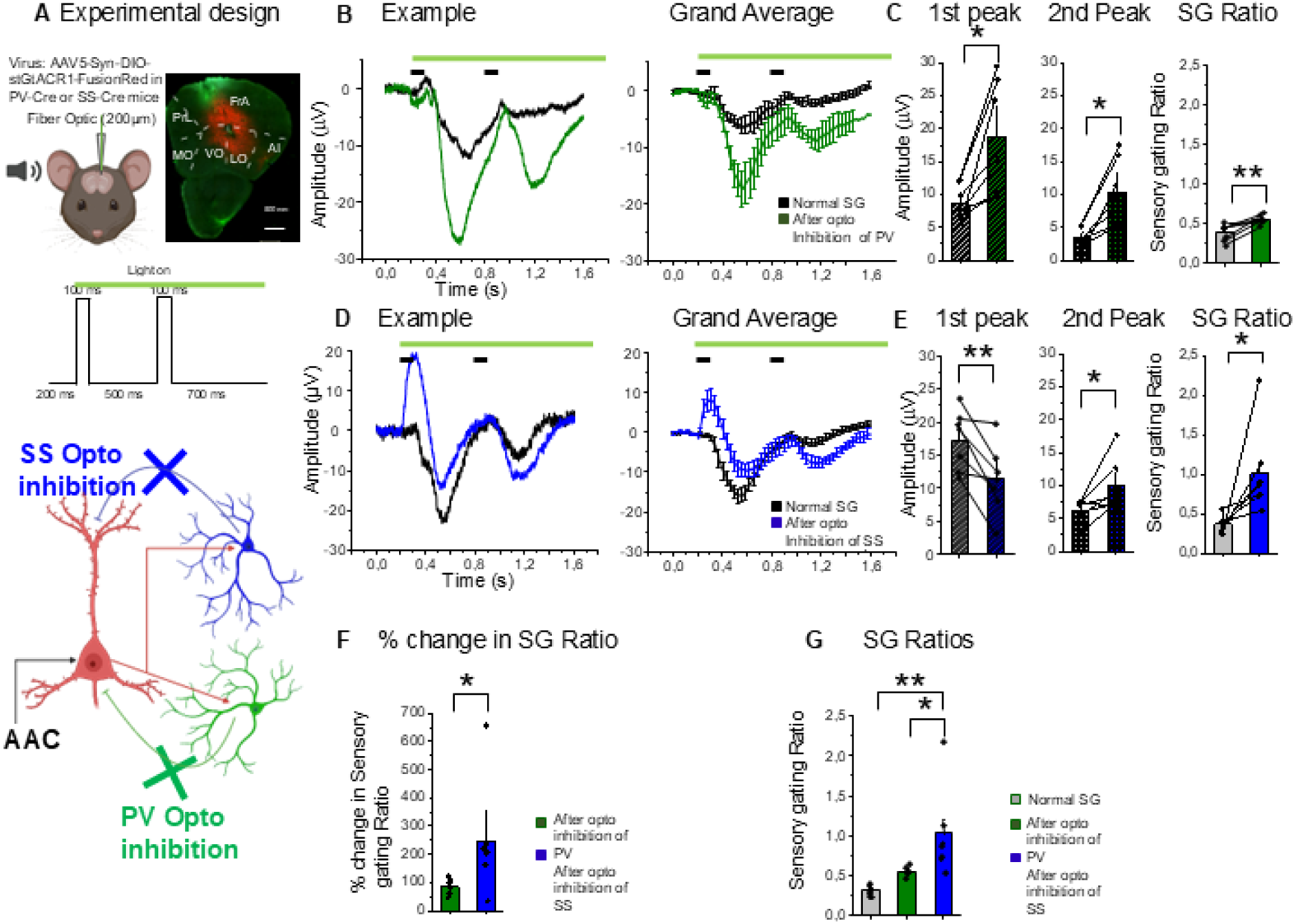
Optogenetic inhibition of SS-interneurons is significantly more effective in counteracting SG compared to PV-interneurons. **A.** Experimental sketch. Optogenetic inhibition of different classes of GABAergic neurons (SS- or PV-interneurons, blue and green, respectively) was performed via the optic fiber inserted in the recording pipette (see Methods). The powerful inhibitory opsin GTCATR1 was used, and the light was on during the SG protocol (green line). **B.** Examples and grand-averages of the LFP responses recorded during the SG protocol with optic fiber off (black) and on (green), when GTCAR1 was expressed in PV interneurons. Note that both responses increased in a quasi-multiplicative way. **C.** Quantifications of the data reported in B: responses to 1^st^ sound (left plot: before optoinhibition: 8,6 ± 1,0 µV *vs* after optoinhibtion: 18,8 ± 3,1µV; paired t-test, *p=0,02), 2^nd^ sound (middle plot: before optoinhibition: 3,4 ± 0,6µV *vs* after optoinhibition: 10,4 ± 1,8µV; paired t-test, *p=0,01) and SG ratio with/without (green/black) illumination (right plot: before optoinhibition: 0,30 ± 0,04 *vs* after optoinhibition: 0,55 ± 0,02; paired t-test, **p=0,003) are shown from left to right. Note that both peaks increased (paired t-tests, *p<0,05) and that we measured a slight but significant increase of the SG ratio (paired t-test, **p<0,01). **D.** Examples and grand-averages of the LFP responses recorded during the SG protocol with light switched off (black) and on (blue), when GTCAR1 was expressed in SS-interneurons. Note that the 1^st^ response diminished, whereas the 2^nd^ response increased. **E.** Quantifications of the data reported in **D**: responses to 1^st^, 2^nd^ sound, and SG ratio with/without illumination are shown from left to right. Note that opposite effects of optogenetic inhibition of SS interneurons on the 1^st^ and 2^nd^ responses (1^st^ peak decrease: 17,3 ± 1,7µV *vs* 11,4 ± 1,9µV, before and after optoinhibition, respectively; paired t-test, **p=0,008 – 2^nd^ peak increase: before optoinhibition: 6,2 ± 0,7µV *vs* after optoinhibition: 9,9 ± 1,5µV; paired t-test, *p=0,03 respectively), which results in an effective SG increase (before optoinhibition: 0,37 ± 0,04 *vs* after optoinhibition: 1,01 ± 0,21; paired t-test, **p=0,03). **F**, Optoinhibtion of SS-interneurons (blue) leads to a larger than percentage increase in SG ratio compared to optoinhibition of PV-interneurons (green; % change of SG ratio in SS-Cre = 246,3 ± 73,0 *vs* PV-Cre = 85,6 ± 9,9;t-test *p=0,04).), **H,** Comparison of SG ratio between controls (without light on in PV-Cre animals, black – same results when without light on SS-Cre mice were used), optoinhibition of PV-(green) and SS-interneurons (blue). Only optoinhibition of SS-interneurons was able to significantly change the SG ratio (normal 0,30 ± 0,04 *vs* SS-Cre= 1,01 ± 0,21 *vs* PVCre = 0,55 ± 0,02; 1 way ANOVA **p= 0,002 with Bonferroni post hoc tests **p = 0,002 (normal *vs* SS); *p= 0,04 (PV *vs* SS); p=0,53 (normal *vs* PV))

In the case of NMDA antagonism, recovery of the 2^nd^ response was not simply linked to suppression of peak responses to 1^st^ sound. We thus wondered whether NMDA receptors could contribute to the sustained character of the acoustic response in OFC (Figure S1A,C). Indeed, NMDA receptor antagonism shortened acoustic response durations (Figure 2F; paired t-test, p=0,02).

As two-photon imaging has shown that sensory-driven, sustained dendritic calcium responses in cortical neurons are largely contributed by NMDA receptors(*23*), we wanted to test whether and how acoustically-driven calcium neuropil responses in OFC became changed upon NMDA antagonism. We measured acoustic-driven neuropil calcium responses via fiberphotometric calcium imaging in Thy1-GCAMP6s mice (strain GP4.3). In these mice the genetically encoded calcium indicator GCAMP6s is under the promoter for excitatory neurons Thy1(*24*) and intracranial fiberphotometric calcium signals reflect largely dendritic transients in pyramidal neurons(*25*). To test the hypothesis that NMDA receptors contribute to sustain auditory-driven dendritic calcium responses in OFC, we compared the acoustic-driven calcium-related photomultiplier signal before and after local injection of the NMDA antagonist AP-5 in OFC in response to a single tone (Figure 2H, see Methods). As shown by the examples and grand averages in Figure 2I(upper plots), NMDA antagonism reduced peak calcium transient compared to saline, vehicle (upper and lower plots with red and blue curves, respectively; see (Figure 2J for peak quantifications; N=5 for NMDA antagonism; paired t-test, p=0,01). Conversely, a slight and non-significant increase was observed upon vehicle (saline solution) injection (Figure 2I – plots with blue traces; Figure 2J N=3; paired t-test, p=0,06), excluding that the signal reduction upon NMDA antagonism was due to time-dependent fluorophore bleaching. We also measured a reduced response duration upon infusion of AP-5 (Figure 2K, top; paired t-test, p=0,004), whereas no changes were observed by infusion of vehicle (Figure 2K, bottom; paired t-test, p=0,486).

We also performed calcium fiberphotometric measurements during the SG protocol (Figure S3A). As evident in the example and grand-average of Figure S3B,C, SG was significantly less pronounced when the neuropil (dendritic) signal was measured compared to electrophysiology (still the 2^nd^ response was smaller compared to the 1^st^, Figure S3D, N=10; paired t-test, p=0,04). The comparison of the response during SG and to stimulus-onset-aligned calcium response to 1^st^ sound only shows that the negligible level of SG was primarily due to the very prolonged character of the calcium transient in response to the first sound (Figure S3C). As a result, SG measured by calcium photometry was statistically smaller compared to the dramatic SG measured with electrophysiology (Supplementary Figure 2D, right; t-test, p=0,000002).

Taken together, our data indicates that local NMDA antagonism in mouse OFC counteracts SG while reducing acoustically-driven dendritic calcium responses.

### Optogenetic inhibition of somatostatine-interneurons counteracted SG more effectively than parvalbumine-interneurons

We next investigated whether the different inhibitory subnetworks play a differential role in mediating SG in OFC, focusing on the two main classes of GABAergic, inhibitory cell types: parvalbumin-positive (PV) interneurons inhibiting pyramidal cell somatic, spike output and the somatostatin-positive (SS) interneurons inhibiting dendritic synaptic inputs. These two cell types respond differently in vivo in the prefrontal cortex during executive function testing(*26, 27*), and their physiology is impaired in animal models of conditions with SG deficits such as schizophrenia(*28, 29*). To selectively inhibit each type of interneurons, we injected an AAV with a floxed version of the inhibitory opsin GTACR1(*20*) in either PV-Cre or SS-Cre transgenic mice, expressing CRE recombinase in PV- or SS-interneurons, respectively (Figure 3A). Electrophysiological recordings were done with the fiber optic cannulated in the electrode holder so to record both pre- and post-illumination responses, and the illumination lasted for the entire SG protocol (both time windows, Figure 3A).

We first photoinhibited PV interneurons (Figure 3B,C) during the SG protocol. As the example and grand-averages of Figure 3B show, photoinhibition of PV-interneurons caused a quasi-multiplicative increase in both 1^st^ and 2^nd^ peak amplitudes. For quantification see Figure 3C: N=7; 1^st^ peak: paired t-test, p=0,02; 2^nd^ peak: paired t-test, p=0,01. The result of the peaks increasing almost in proportion was that the SG ratio was only slightly affected by optoinhibition of PV-interneurons, although the difference was significant as the increase of the 2^nd^ response was slightly but significantly larger compared to the increase of the 1^st^ response (Figure 3C, paired t-test, p=0,003).

The result of SS-interneuron optoinhibition, done in the same conditions, were qualitatively different. The upward LFP signal deflection we saw at illumination onset reflects an opsin-activation related signal, as this upward deflection was also present when the optic fiber was switched on in absence of sound (see black grand-average of Figure S4A). This upward deflection could be generated by a light-driven inhibition (as locally-originated upward LFP signals represent a local inhibitory source(*30*)), possibly due to the fact that SS-interneurons powerfully inhibit PV-interneurons(*31*). In line with the idea that the initial light-driven upward signal is at least partially of GABAergic nature, we observed that extracellular application of GABA antagonists reduced this peak (see Figure S4A, red grand-average, N=3). The examples and grand-averages of Figure 3D show that SS-interneurons photoinhibition during SG reduced the1^st^ sound downward response while increasing the 2^nd^ response (Figure 3E: 1^st^ peak: paired t-test, p=0,008; 2^nd^ peak: paired t-test, p=0,03). Therefore, there was a statistically significant increase in the SG ratio before and after SS-interneuron photo-inhibition (Figure 3E, paired t-test, p=0,03). We also found that it was enough to optoinhibit SS-interneurons during the 1^st^ sound time window to achieve unmasking of the 2^nd^ response (Figure S5).

Thus, optoinhibition of SS-interneurons was about 3 times effective in counteracting SG compared to PV-interneurons as shown by a change of SG ratio of 246% vs 85% (on average values; Figure 3F; t-tests, p=0,04). Unpaired comparison of SG ratios among controls (SG before optoinhibition in PV-Cre animals), inhibition of PV-interneurons and SS-interneurons, showed that only optoinhibition of SS-interneurons was able to significantly counteract SG compared to both control and optoinhibition of PV-interneurons (Figure 3G; One way ANOVA, p = 0,002, with *post-hoc* Bonferroni tests; p=0,002 (normal *vs* SS); p=0,04 (PV *vs* SS); p=0,53 (normal *vs* PV)).

### The results of optogenetic manipulations are accounted for by a phenomenological rate model requiring longer temporal integration of SS-interneurons compared to PV-interneurons

We constructed and trained a phenomenological firing rate model to reproduce the experimental LFP curves to reveal potential underlying circuit properties that could account for the observed curve shapes and gating ratios. We explicitly modeled firing rates for populations of pyramidal cells as well as three distinct interneuron subtypes: SST, PV and VIP cells, and modeled their mutual influences according to a simplified but biologically plausible topology at the population level, previously studied in (Figure 4A) (*32*). Sound pulses and optogenetic currents were modeled as time-dependent inputs with fixed parameters, acting on the corresponding targets (the AAC population for sound, and the appropriate interneuron population for each optogenetic intervention; see methods for details), but the strengths of the interactions between populations, as well as the intrinsic time constants for the evolution of each population’s firing rate, were left as free parameters.

**Figure 4.**
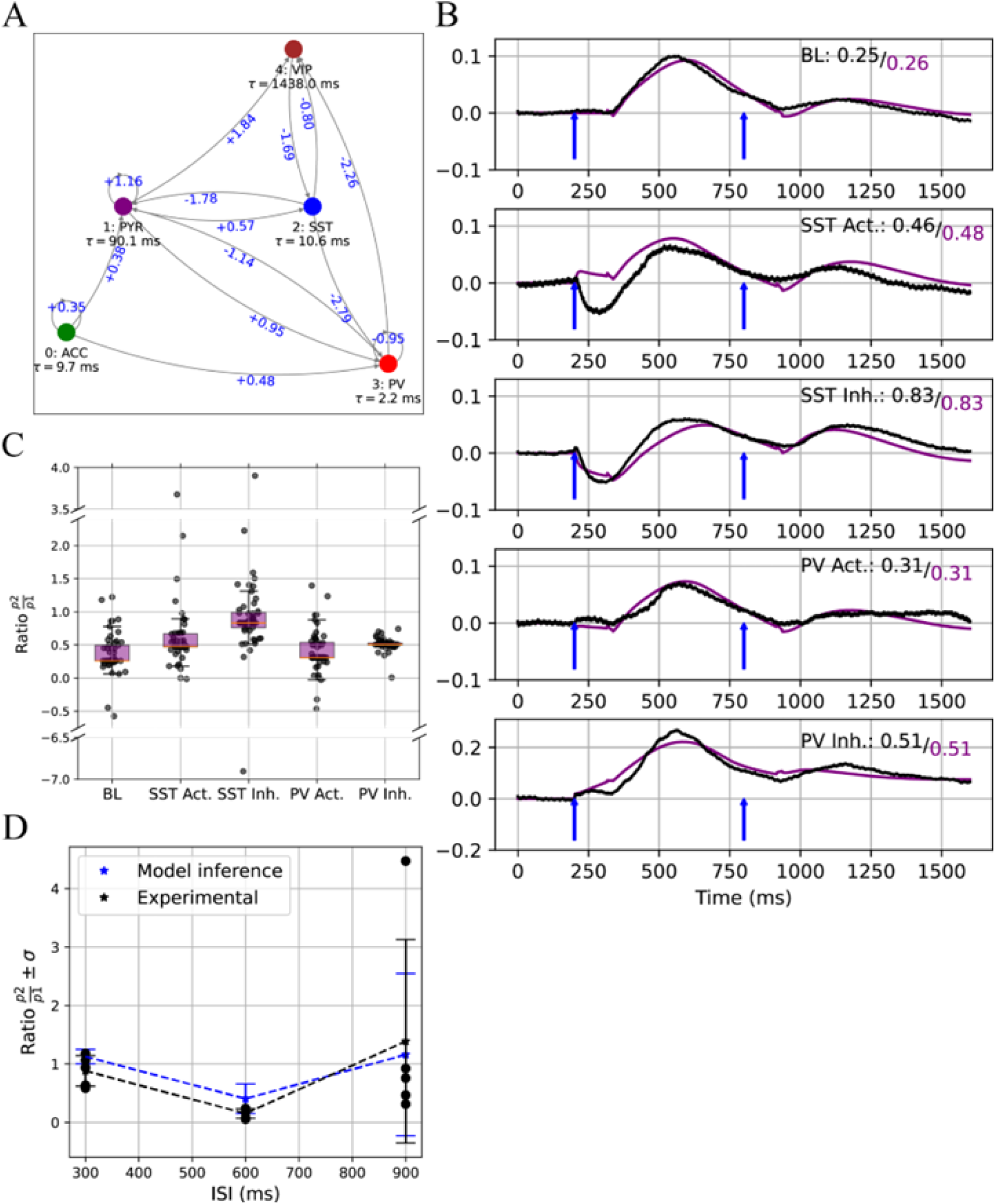
A. Simulated topology and corresponding interaction strengths and intrinsic time-constants for a phenomenological, biologically plausible model of excitatory and inhibitory circuitry in prefrontal auditory gating. Connection strengths and intrinsic time-constants were determined by optimizing the similarity between experimental LFPs and total inputs at each node (see *Methods*), while the topology was held fixed. Note the different time-constants for the SST and PV nodes, a key feature necessary for the model to reproduce experimental observations. **B.** Time-dependent input curves and corresponding observed LFPs for five distinct experimental conditions; top to bottom: baseline, optogenetic excitation of SST cells, optogenetic inhibition of SST cells, optogenetic excitation of PV cells, optogenetic inhibition of PV cells. Purple: simulated time-dependent sum of inputs at the PYR node. Black: negated experimental OFC LFPs. Inlaid texts show resulting gating ratios for experimental (black) and simulated (purple) curves corresponding to each condition. Time-dependent inputs and outputs for all nodes and conditions are shown in supplementary materials (Figure S6). **C.** Effect of perturbing individual model parameters on the simulated gating ratios, for each condition. Each black circle corresponds to an individual run, where a single parameter was perturbed either +5% or −5% away from the value that results in the optimal similarity between simulated curves and experimental LFPs. The corresponding curves are shown in supplementary materials (Figure S7). **D.** Comparison between blind model inferences and experimental observations for additional experiments where the inter-stimulus intervals (ISI) were changed. Model parameters were optimized considering exclusively the experiments with *ISI* = 600*ms*; results for different ISI values were not known at the time of model training. Black dots represent individual experiments; distributions for the model inference results were generated by introducing a low level of additive white noise at the PYR node and running 50 distinct simulations for each ISI value (individual runs not shown). Note that the model correctly predicts the unseen experimental results, reproducing the observation that the peak gating happens at *ISI* = 600*ms* as well as the return to baseline and eventual observations of anti-gating at long ISI intervals. Experimental data: Interstimulus time dependence of SG. 500ms evoked the strongest SG (200ms: 0,80 ± 0,12; 500ms: 0,15 ± 0,04; 800ms: 0,61 ± 0,14; 1100ms: 0,87 ± 0,1; 1 way ANOVA RM *p= 0,002 with Bonferroni post hoc tests: 500 vs 200, *p=0,02; 500 vs 800 p=0,12, *500 vs 1100 p=0,01).

All free parameters were in turn determined via optimization of the similarity between experimental LFPs measured at the AAC and OFC, and the time-dependent inputs at the corresponding AAC and PYR populations, simultaneously for all experimental conditions. This choice is consistent with an interpretation of LFPs as being driven by subthreshold dendritic currents at the measurement site(*33*), and not necessarily correlated to the eventual spiking output at the same site. After a small number of rounds of optimization, the simulated population activities reproduced the experimental curve shapes and gating ratios with great fidelity (within-sample mean squared error MSE = 3.36⨯10-4; Figure 4B,C). Note how the ca 5-fold longer time constant predicted by the model for SS-interneurons as compared to the one of PV-interneurons, indicative of a longer time integration window of SS-interneurons.

We further tested the obtained solution by simulating the model subject to a novel condition – the presentation of pairs of pulses under different inter-stimulus intervals not present in the training dataset – and comparing the model predictions to a new set of unseen experimental data. The model was able to qualitatively reproduce these unseen results with surprising accuracy (Figure 4D), suggesting that this simplified approach was able to capture features of the underlying biological microcircuit involved in SG.

### Blockade of dopaminergic modulation by neuroleptics reduces acoustic responses in both AAC and OFC but affects the SG ratio only in OFC

Unbalances in dopaminergic neuromodulation have been associated with all aspects of schizophrenia and other psychotic disorders, with the dominant model positing a reduced dopaminergic transmission in medio-orbital prefrontal cortex mediating negative and cognitive (dysexecutive) symptoms. We thus investigated the effect of dopaminergic transmission blockade by comparing SG before and 30min after a local intracortical injection of the typical neuroleptic flupentixole (300nL, 5 mM as in(*34*)), effectively antagonizing both D1 and D2 receptors (Figure 5A). As shown by the examples and grand-averages of Figure 5A,B, we consistently observed a strong depression of 1^st^ response (Figure 5C; N=6; paired t-test, p=0,02) accompanied by an increase of the 2^nd^ (paired t-test, p=0,03), ruling out a generalized divisive downscaling of responses. Consequently, the SG ratios became significantly higher after flupentixole injection (Figure 5C, paired t-test, p=0,03). We also measured the neuroleptic capacity to affect response duration, by comparing the half width of the response to 1^st^ sound before and after flupentixole injection. Neuroleptic treatment reduced acoustic response duration in OFC (Figure 5D, paired t-test, p=0,03).

**Figure 5.**
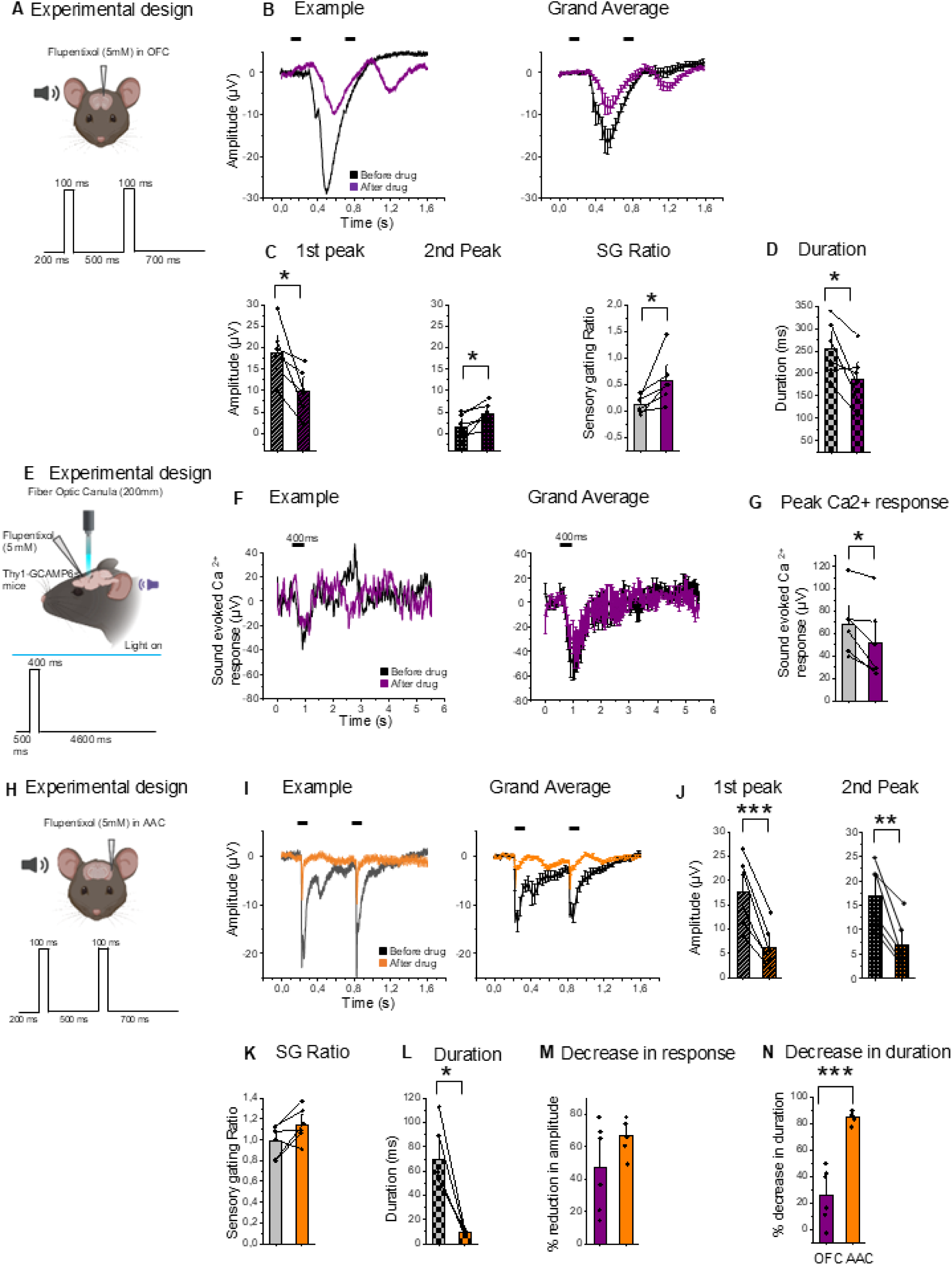
Local infusion of typical neuroleptics counteracts SG in a way qualitatively similar to optoinhibiton of SS-interneurons, while reducing sound-evoked calcium responses. **A.** Experiment sketch: the typical neuroleptic flupentixol was injected (0,3μL, 5mM) through the recording pipette to compare SG before and after drug application in OFC. **B.** Examples and grand averages of LFP responses during SG protocol before and after drug application (black and magenta, respectively). **C.** Note the reduction of the 1^st^ response (left plot: before drug 18,9 ± 2,6µV *vs* after drug: 9,9 ± 2,1µV; paired t-test, *p=0,02) and the increase of the 2^nd^ response (middle; before drug: 1,6 ± 1,1µV *vs* after drug: 4,6 ± 1,1µV; paired t-test, *p=0,03), which caused a significant increase of the SG ratio (right plot; before drug: 0,12 ± 0,06 *vs* after drug: 0,58 ± 0,19; paired t-test, *p=0,03). **D.** Neuroleptics reduced slightly but significantly the duration of the first acoustic response in OFC (255,2 ± 25,3ms *vs* 186,7 ± 25,3ms; paired t-test, *p=0,03). **E.** Experiment sketch of the calcium photometry in Thy-1 GCAMP6s mice before and after neuroleptic injection. Injection of the neuroleptic was done through an injection pipette inserted in a neighboring craniotomy. **F.** Examples and grand averages of calcium-related responses before and after flupentixole injection (black and magenta, respectively). **G.** The peak response was reduced upon neuroleptic injections (before drug: 68,0 ± 11.3µV *vs* after drug: 52,1 ± 13.6µV; paired t-test, *p=0,017). **H.** Same experiment in **A** but performed in acoustic cortex (AAC). **I.** Examples and grand averages of LFP responses during SG protocol before and after drug application (black and orange, respectively). Observe a dramatic reduction of both acoustic responses. **J,K.** Quantifications of the neuroleptic-driven response reduction for the 1^st^ and 2^nd^ peak (1^st^ response, left: before drug: 17,5 ± 2,9µV *vs* after drug: 6,2 ± 1,7µV; paired t-test, ***p=0,0005; 2^nd^ response, middle: before drug: 16,9 ± 2,6µV *vs* after drug: 7,0 ± 2,0µV; paired t-test, **p=0,003), so that the SG ratio was not affected (panel K, right; before drug: 0,99 ± 0,06 *vs* after drug: 1,14 ± 0,06; paired t-test, p=0,07). **L.** Neuroleptics reduce the duration of the first acoustic response in AAC (before drug: 70,0 ± 10,3 *vs* after drug: 9,8 ± 1,0ms; paired t-test, **p=0,002). **M,N.** Neuroleptic induced similar percentage changes in response amplitude in OFC and AAC (panel **M**; OFC: 47,5 ± 11,0 *vs* AAC: 67,1 ± 4,4; t-test, p=0,13) but were more effective in reducing the duration of acoustic responses in AAC (orange) vs OFC (magenta) (panel **N**; right: OFC: 26,0 ± 8,5 *vs* AAC: 85,2 ± 1,90; t-test, ***p=0,00004).

The capacity of the neuroleptic flupentixole to reduce response duration was reminiscent of the one observed after NMDA blockade. As we showed that NMDA receptors are a major determinant of sound-evoked calcium dendritic response, we wanted to test whether neuroleptic treatment was also able to reduce sound-evoked neuropil calcium responses measured by fiberphotometry in GCAMP-Thy1 mice (Figure 5E). The results showed that injection of flupentixol caused a slight but significant negative modulation of sound-evoked neuropil calcium response (Figure 5F,G, N=6; calcium related PMT signal: paired t-test, p=0,017).

As dopaminergic innervation is significantly denser in OFC compared to more caudal sensory areas such as AAC(*35*), we compared the effects of flupentixole injection in AAC (experiment sketch Figure 5H), the main input of acoustic responses in OFC(*17*). As shown in Figure 5I, flupentixole reduced dramatically sensory responses in AAC to *both* sound responses (Figure 5J, N=6; 1^st^ response, paired t-test, p=0,0005; 2^nd^ response, paired t-test, p=0,003), without affecting the SG ratio (Figure 5K: paired t-test, p=0,07). Neuroleptic treatment reduced the acoustic response duration significantly in AAC as well (Figure 5L, paired t-test, p=0,002).

To compare neuroleptic effects on sensory responsiveness *per se* in the two areas, we measured the percentage reduction in response amplitude and duration caused by neuroleptics in AAC and OFC to the 1^st^ sound response (as the two responses were similar in AAC). The percentage reduction of response amplitude was not statistically different in AAC compared to OFC (Figure 5M; t-test, p=0,13). The flupentixole-induced percentage decrease in response duration was significantly more pronounced in AAC compared to OFC (Figure 5N, t-test, p=0,00004). Thus, neuroleptics reduced acoustic response duration proportionately more in AAC and OFC.

Taken together, these data show that local antagonism with dopaminergic transmission in OFC effectively counteracts SG, in a way similar to the optogenetic inhibition of SS-neurons (reduction of 1^st^ peak, increase of 2^nd^). This effect is accompanied by a negative modulation of neuropil, dendritic calcium sound-evoked calcium response and is not attributable to a generalized suppression of responsiveness, as the 2^nd^ responses increases in OFC on one side, and as both acoustic responses were also reduced in AAC without an increase of the 2^nd^ response in AAC on the other side.

### Neuroleptic effects on SG are reduced by optogenetic activation of somatostatin-positive, but not of parvalbumin-positive interneurons

As neuroleptics counteracted SG in a way comparable to photoinhibition of SS-interneurons (decreasing the 1^st^ peak and increasing the 2^nd^), we tested the hypothesis that neuroleptics effect could be at least in part due to inhibition of SS-interneurons, in line with the idea that dopamine positively modulates SG via SS-interneurons(*36*). To test this hypothesis, we measured whether neuroleptic effects on SG could be at least partially counteracted by optogenetic stimulation of SS-interneurons. We injected in OFC AAVs containing a floxed channelrhodopsin (ChR2) sequence either in SS-Cre or in PV-Cre mice, so to express the excitatory opsin in either population of inhibitory cells. We first controlled that optogenetic activation of inhibitory interneurons reduced acoustic responses, by recording with the optic fiber in the recording pipette (Figure S8A). Optoactivation of both PV- or SS-interneurons reduced the amplitude of the 1^st^ peak (Figure S8; panels B,C for SS-Cre mice, N=6; 1^st^ peak: paired t-test, p=0,02; panels D,E for PV-Cre mice, N=4; 1^st^ peak: paired t-test, p=0,01), without changes of the already suppressed 2^nd^ peak amplitude (SS-Cre mice: paired t-test; p=0,96; PV-Cre mice: paired t-test, p=0,96). The resulting effects on the SG ratio were not significant (Figures S8C,E; SS-Cre paired t-test, p=0,15; PV-Cre paired t-test, p=0,66).

We next compared SG upon neuroleptic with/without photoactivation of the two types of GABAergic interneurons Figure 6A). As the examples and grand averages of Figure 6B show, SS-interneurons photoactivation resulted ina kind of subtractive effect that however did not significantly affect the first response, whereas it caused a consistent and significant decrease of the 2^nd^ response (Figure 6C for quantification; 1^st^ paired t-test, p=0,36; 2^nd^ paired t-test, p=0,02). ChR2 activation in SS-interneurons significantly reduced the higher-than-normal SG measured upon neuroleptic infusion (Figure 6C, paired t-test, p=0,02), that is neuroleptic effect on SG was significantly counteracted by SS-interneurons photostimulation. This effect was cell type-specific as PV-interneurons stimulation did not affect any of the peaks (Figure 6D, E; 1^st^ peak, paired t-test, p= 0,39; 2^nd^ peak, paired t-test, p=0,9). Therefore, the higher-than-normal SG ratio induced by NL was not diminished by photostimulation of PV-interneurons (Figure 6E; paired t-test, p=0,2).

**Figure 6.**
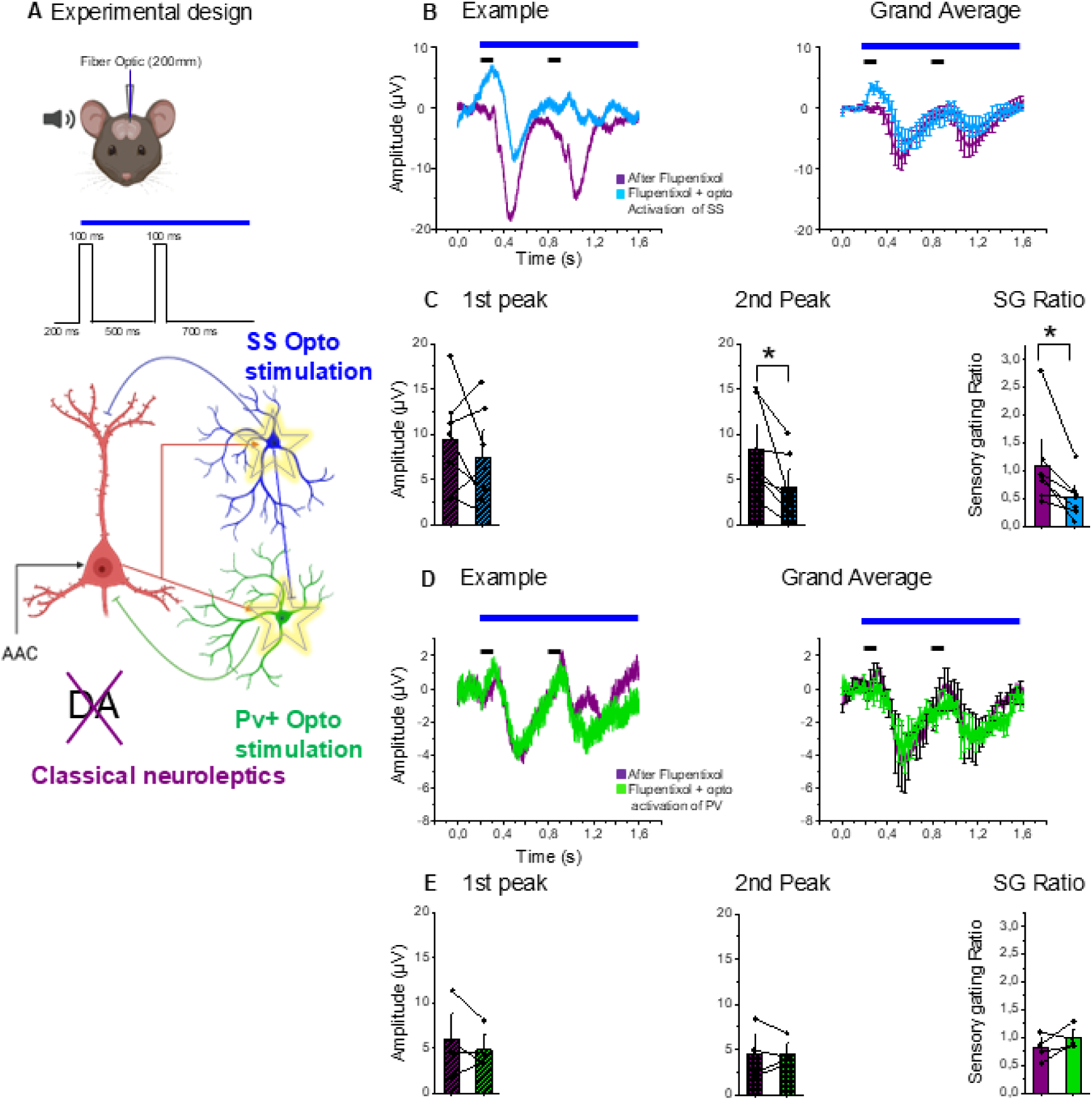
Neuroleptic effect on SG is significantly counteracted by optogenetic activation of SS-but not of PV-interneurons. **A.** Experiment sketch. The recording pipette that infused flupentixol (0,6μl, 5mM) contained a cannulated optic fiber to optogenetically inhibit either PV- or SS-interneurons during the SG protocol. We compared the neuroleptic effect on SG before and during optostimulation of the two types of GABAergic interneurons. **B.** Photostimulation of SS-interneurons effectively counteracted neuroleptic effect on SG. Examples and grand averages of SG upon neuroleptic application with the optic fiber off and on (magenta and light blue, respectively). Note the difference in the SG ratio, due to a visible subtractive effect that had a larger relative impact on the 2^nd^ response. **C.** Quantification of data from the above-mentioned experiment. SS-interneuron photostimulation reduced the SG ratio by selectively reducing the 2^nd^ response (1^st^ response – left plot: before *vs* after photoactivation: 9,3 ± 2,1µV *vs* 7,4 ± 2,0µV; paired t-test, p=0,36; 2^nd^ response – middle plot: before vs after photoactivation: 8,3 ± 1,8µV *vs* 4,0 ± 1,4µV; paired t-test, *p=0,02; SG ratio reduction– right plot: before vs after photoactivation: 1,09 ± 0,30 *vs* 0,52 ± 0,14; paired t-test, *p=0,02). **D.** Photostimulation of PV-interneurons did not affect the SG ratio in neuroleptic-injected mice. Examples and grand-averages in mice injected with neuroleptics with optogenetic stimulation off or on (magenta and green, respectively). Note the lack of effect of PV-interneuron photostimulation on SG. **E.** Photostimulation of PV-interneurons did not change 1^st^ response, 2nd response nor the SG ratio (1^st^ peak - left plot: before and after photoactivation of PV-interneurons: 5,8 ± 2,0µV *vs* 4,8 ± 1,1µV; paired t-test, p=0,39; 2^nd^ peak - middle plot: before and after photoactivation of PV-interneurons: 4,5 ± 1,5µV *vs* 4,5 ± 0,8µV; paired t-test, p=0,9; SG ratio-right plot: before vs after photoactivation: 0,81 ± 0,12 vs 1,00 ± 0,10; paired t-test, p=0,2 respectively).

Taken together, these data show that photoactivation of SS-interneurons, but not of PV-interneurons, significantly counteracted the SG deficit induced by acute, local blockade of dopaminergic transmission in OFC.

## DISCUSSION

### Synaptic mechanism mediating cell-type specific role of GABAergic cells in SG: subtractive vs divisive inhibition

Three observations indicate that the inhibitory gating we documented happens mostly at dendritic input, rather than at somatic output level. First, inhibiting dendrite-targeting, SS-interneurons affects SG more effectively compared to inhibition of soma-targeting, PV-interneurons. Second, manipulation of GABA_B_ receptors, which are mostly localized in the dendritic tree and not at somatic level, contributes significantly to modulate SG. However, the possibility that GABA_B_ receptors could also affect axonal release of glutamate from AAC axons exists, as GABA_B_ receptors are expressed at presynaptic level(*37*): even in this case, SG would actually happen across the synapse AAC-OFC. Third, the fact that NMDA receptor antagonism reduced neuropil calcium response, largely of dendritic origin in Thy1 GCAMP6 mice(*24*), and also affects SG, is in line with the idea that NMDA-dependent, dendritic calcium influx is needed for SG. Our finding that SS-interneurons are key cellular mediators of SG is in line with literature showing that SS-interneurons mediate subtractive forms of inhibition(*38, 39*), whereas PV-interneurons would instead mostly mediate divisive, or normalizing inhibition, which scales proportionately responses (but see (*40*)). Indeed, input subtraction, but not division, is by definition able to modify the SG ratio (result of division between the two response amplitudes). As far as SS-interneurons are concerned, the effects of their optogenetic manipulations are the result of a complex network effect, particularly *in vivo*, as optogenetic manipulation of one cell type by necessity reverberates on all the other connected ones. In particular, as SS-interneurons inhibit PV-interneurons, PV-interneurons would be disinhibited upon inhibition of SS-interneurons. this PV-interneurons disinhibition could thus account for the upward initial deflection upon SS-interneurons optoinhibition on one side, and for the fact that this very same manipulation reduced the 1^st^ response. Unmasking of the 2^nd^ response upon SS-interneurons inhibition is more probably mediated by direct blockade of dendritic inhibition, mediated to a significant extent by GABA_B_ receptors, as a sustained PV-interneuron disinhibition would have further decreased the 2^nd^ peak instead of increasing it. Similarly, the result of pharmacological GABAergic blockade showing that the 1^st^ peak was unchanged could be accounted for the fact that we were simultaneously blocking postsynaptically the GABA released by both PV-interneurons (whose blockade should have increased the 1^st^ response) and by SS-interneurons (whose blockade should have decreased it), thus eliciting two mutually cancelling responses.

### Dopaminergic modulation of GABAergic interneuron function

The classical dopaminergic hypothesis of schizophrenia postulates a reduction of dopaminergic transmission at prefrontal cortical level. Local infusion of the 1^st^ generation neuroleptic flupentixole counteracted SG by reducing the 1^st^ peak and increasing the 2^nd^ one, thus altering the normal SG ratio. Manipulation of dopaminergic transmission is known to affect SG(*5*). However, the effects of typical neuroleptics are controversial, although blockade of SG has been reported in rats(*12, 41*). Our data, together with the observation that increase of dopaminergic transmission (e.g. via amphetamine) also counteracts SG, suggest that perhaps both blockade and great increase of dopaminergic transmission counteracts prefrontal SG. This is reminiscent of the U-shaped curve of dopamine effect on working memory(*42*), where too high or too little dopaminergic transmission at prefrontal level equally disrupts working memory. Possibly, for both working memory and SG, two fundamental prefrontal executive functions, an ideal middle range of dopaminergic transmission is required.

Flupentixole effectively counteracts OFC SG in a way quite similar to optoinhibition of SS-interneurons (reduction of 1^st^ response and increase of the 2^nd^), raising the possibility that neuroleptics effects were at least partly mediated by SS-interneurons inhibition. Dopamine effects on prefrontal inhibitory microcircuits are complex and layer-specific(*43*), with literature converging on the fact that dopamine increases PV-interneurons excitability(*44, 45*).

Less is known on the action of dopamine on SS-interneurons, although a facilitatory action has been hypothesized(*36*). Our observation that neuroleptic effects are at least partially counteracted by photostimulation of SS-, but not of PV-interneurons, is in line with the idea that dopamine could facilitate SG via SS-interneurons activation. Of relevance, the disinhibitory effects of acute dopaminergic blockade on dendritic activity *in-vivo* in motor cortex can be overcome by chemogenetic activation of SS-interneurons in mouse motor cortex(*46*). SS-interneuron photostimulation reduced the 2^nd^ response only upon neuroleptic infusion but not in controls, in line with the idea that in controls the inhibitory gating mechanism is already activated at quasi-saturating level, whereas upon neuroleptic application, SS-interneurons and their effects can be reactivated optogenetically as they were partially blocked by neuroleptics. Future studies are needed to better investigate the prefrontal effect of dopaminergic stimulation in cell-type-specific subnetworks.

### A microcircuit mechanism mediating prefrontal SG

Our results allow us to sketch a model of SG microcircuit basis. First, reducing pyramidal neuron activity during the 1^st^ response reinstates a significant response to the 2^nd^sound, indicating that pyramidal neurons activity in response to 1^st^ sound recruit SS-interneurons. This activation is more sustained in OFC compared to its input area AAC. Of relevance, prefrontal neurons have an intrinsic tendency to fire in a prolonged way, as shown by the “sustained firing” during working memory tasks(*47–49*). Interestingly, both NMDA receptors(*50*)and dopamine(*51*), whose interference reduces SG, greatly facilitate prefrontal neuron persistent activation (as well as working memory), and – as our data show - also the duration of OFC sound-driven electrophysiological activation. Our calcium photometric data from Thy-1 animals reflect mostly calcium dendritic activity of pyramidal dendrites(*24*) and show a strong reduction of sound-driven calcium signal upon NMDA receptor blockade, in terms of both amplitude and duration, in line with the idea that NMDA-evoked increases of dendritic calcium could be a mechanism behind the sustained duration of OFC electrophysiological acoustic responses. NMDA-driven dendritic calcium spikes indeed propagate to and prolong somatic activation(*52*), contributing to sharpen response selectivity of pyramidal neurons(*23, 53*). Thus, the sustained character of 1^st^ sound-evoked response typical of OFC, determined at least partially by NMDA-dependent calcium, is plausibly needed for SS-interneurons activation at threshold. Paired recordings have indeed shown that whereas pyramidal-to-PV-interneuron synapses are depressing during pyramidal neurons sustained firing, pyramidal-to-SS-interneuron synapses potentiate their response by temporal summation, both *in-vitro*(*54*), *in-vivo*(*55*), and also in living human brain slices(*56*). Critically, this potentiation happens because SS-interneurons express a subunit of NMDA receptors that prolongs unitary excitatory postsynaptic potentials duration(*57, 58*) whereas PV-interneurons loose this isoform at adolescence(*59*). NMDA receptors could thus facilitate SG by sustaining pyramidal neurons firing on one side, but also by facilitating input summation in SS-interneurons dendrites on the other side (see Figure 7). Indeed, SS-interneurons would be recruited during the 1^st^ sound response, as confirmed by the fact that optoinhibition of SS-interneuron inhibition during the 1^st^ time window is enough to reinstate 2^nd^ sound responses (Figure S3). The circuit model we propose accounts for the fact that knocking out NR2B receptors selectively from SS-interneurons in medio-orbital cortex induces a series of behavioral deficits including reduced paired pulse inhibition(*60*), the motor read-out of SG (although electrophysiological SG and behavioral paired-pulsed inhibition do not necessarily correlate(*61*)). Noticeably, our data are accounted by a connectivity model requiring SS-interneurons to have 5-fold longer temporal dynamics compared to PV-interneurons, a ratio consistent (within a factor of 2) with the one found in connectivity studies in vivo by Petersen and colleagues (Figure 3 in (*55*)). Our findings that SS-interneurons and GABA_B_ receptors are key cellular and molecular players behind prefrontal SG respectively, are in line with the fact that SS-interneurons modulate cortical activity via GABA_B_ receptors(*62*). The time frame of action of GABA_B_ receptors matches with the optimal interstimulus time for SG (see inference part in our modelling work), as it takes between 150-400ms for GABA_B_ to exert its action(*30*).

**Figure 7.**
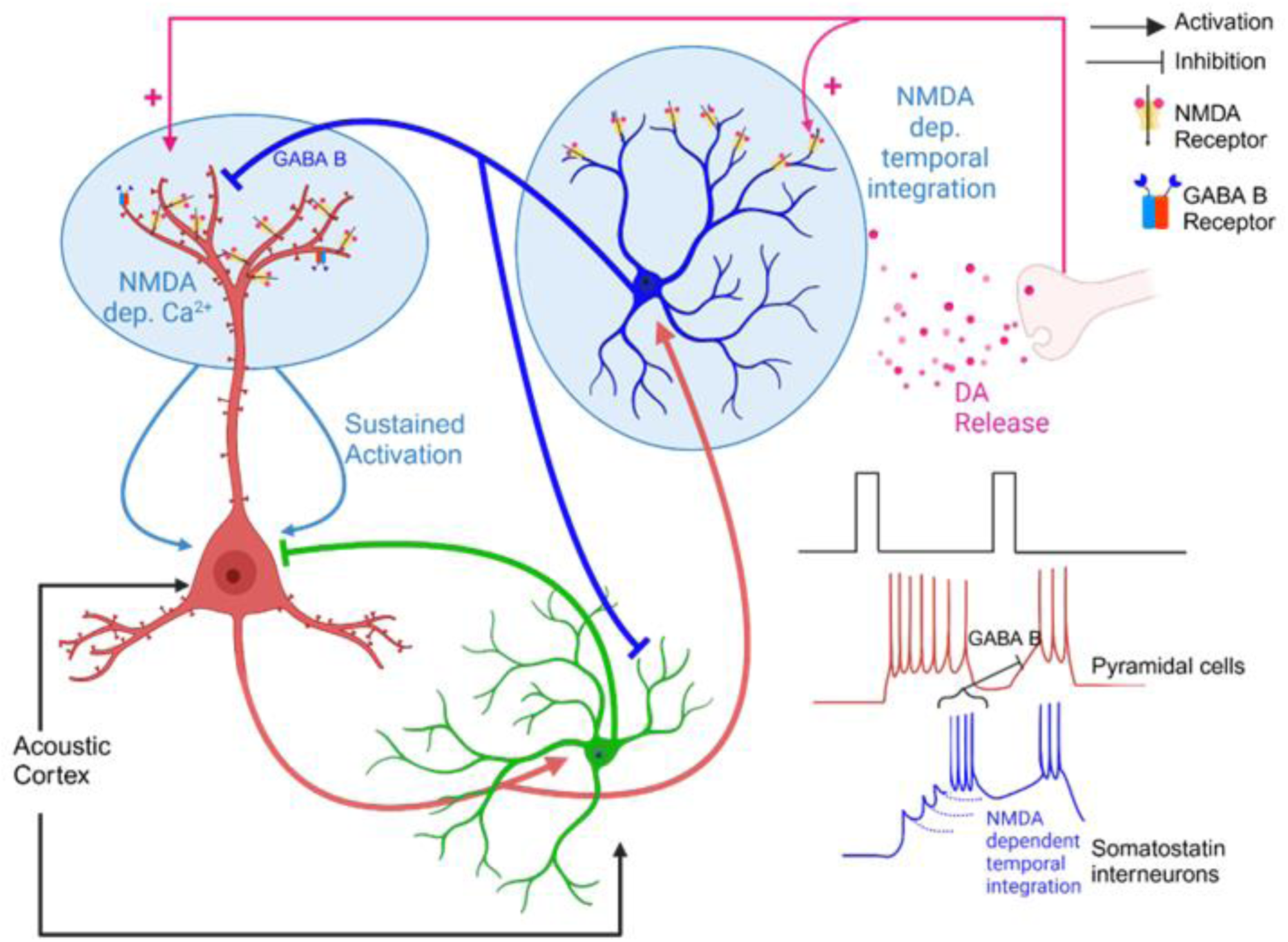
A model microcircuit behind prefrontal SG. Activation of OFC pyramidal neurons from AAC gives rise to a sustained synaptic response, which is facilitated by dendritic NMDA receptor activation on one side, and by the strong prefrontal dopaminergic neuromodulation on the other side. Somatostatin-interneurons are better suited as temporal integrators of the pyramidal sustained activity, due to their longer NMDA unitary postsynaptic potentials, compared to PV-interneurons(*55*). The progressive recruitment of SS-interneurons in response to the first excitatory response might bring them to threshold so that the released GABA can silence the second response, within a time frame compatible with recruitment of dendritic GABA_B_ receptors(*30*). Dopamine facilitation of SG might be due to dopamine capacity to enhance sustained activation, possibly via facilitation of NMDA receptors action(*64*), in the dendrites of both pyramidal and SS-interneurons. Note the crucial role of NMDA dendritic receptors in the sustained pyramidal neuron response on one side, and on promoting temporal summation in SS interneurons dendrites. The strong dopaminergic innervation of the prefrontal areas and the selective expression of NR2B subunits in SS interneurons in prefrontal areas are the documented area-specific mechanisms proposed to be behind the prefrontal specific SG, at the light of the findings of this experimental work.

Our results show that dopaminergic neuromodulation facilitates SG (as its blockade via typical neuroleptics disrupts it). This facilitatory action could happen as dopamine can facilitate NMDA receptor function on one side(*63, 64*) which in turn could support sustained activation of prefrontal neurons (e.g. during working memory), but also via facilitation of NMDA-dependent integration in SS-interneurons (see model microcircuit Figure 7). Moreover, acute dopamine blockade increases pyramidal neuron dendritic activity, an effect overcome by chemogenetic activation of SS-interneurons(*46*), in line with our results that dopaminergic blockade can be overcome by optogenetic stimulation of SS-interneurons.

### Cortical area specificity of SG

We found a dramatic difference in SG between OFC and its main cortical input AAC(*17*), which suggested the role of a local, prefrontal inhibitory mechanism. Similarly, SG is very pronounced in the dorso-lateral frontal association area, but not in sensory association areas such as the PPC, that also sends axons to OFC(*19*). SG can be measured in rodent hippocampus(*12*) or even primary auditory cortex or brain stem nuclei(*11*). SG in prefrontal targets like hippocampus could be explained by a postsynaptic phenomenon compared to prefrontal cortex, whereas the way smaller SG in acoustic cortex(*11*), may a sensorimotor paired-pulsed inhibition component in the awake animal. Of relevance, motor- and acoustic-related modulation of the membrane potential even in visual cortex has been reported(*65*). Undoubtfully, SG in auditory cortex(*11*) is way smaller (negligible in our measurements) compared to SG in OFC. Our data indicates that prefrontal areas have hardwired synaptic machinery performing SG even in the anaesthetized, unconscious brain state.

What could be the mechanisms accounting for the area-(prefrontal) specificity of SG? The persistent firing (or sustained activation at subthreshold/synaptic input level) is a feature of prefrontal neurons favored by both NMDA receptors(*50*) and dopamine receptors(*51*). Moreover, dopamine can greatly enhance NMDA currents in prefrontal cortex (e.g.(*64, 66*). Prefrontal cortical neurons begin to express NR2B isoform of NMDA receptors (facilitating both persistent firing and temporal summation due to the longer tau-off of excitatory postsynaptic potentials) at a time of adolescence(*59*). Importantly, this developmental switch at adolescence is not happening in other cortical areas such as the visual cortex(*67*), indicating the emergence of an area-specific mechanism facilitating persistent firing. Moreover, this molecular switch is cell type-specific, as it happens in SS-interneurons but not in PV-interneurons, and hence it promotes input temporal integration selectively in SS-interneurons. Another prefrontal area-specific factor is dopaminergic modulation, which shows a higher density of dopaminergic fibers in prefrontal areas compared the rest of neocortex. Our finding that neuroleptics reduce relatively more the duration and amplitude of the acoustic response in AAC compared to OFC is in line with dopamine innervation being stronger in OFC because the neuroleptic antagonism is based on competitive displacement of dopamine from the binding sites (neuroleptics have a harder time antagonizing dopamine where its concentration is higher). Dopamine may boost sustained activation by facilitating NMDA-driven calcium current, as also suggested by our observation that neuroleptics reduces sound-evoked, dendritic calcium transients that we showed to be largely NMDA-dependent. Also, D1 and NMDA receptors co-localize in prefrontal dendritic excitatory spines and synergistically increase intracellular calcium responses(*64*).

Thus, our data support the view that higher prefrontal dopamine innervation and cell-type specific changes in molecular isoform of NMDA receptors occurring in prefrontal SS-but not in PV-interneurons interact at the level of both the pyramidal and SS-interneurons dendrites, so to account for the prefrontal area specificity of the SG we documented.

### Clinical relevance

Schizophrenia is linked and possibly caused by disturbances of the prefrontal input processing circuitry (either externally or internally generated, thus involving both perceptions or thoughts, respectively) and that also controls executive functions such as attentional filtering. Thus, SG deficit in psychosis-probe disorders like schizophrenia represents the biological, endophenotypic expression of an inhibitory circuit vulnerability linked to disturbances of sensory perception such as hallucinations, of thought disorders such as delusions on one side, but possibly also of executive dysfunctions such as attentional problems. Thus, the identification of the cell type specific, dopamine-sensitive, inhibitory prefrontal microcircuit mediating SG restricts and isolates allows to test which mechanistic steps possibly are impaired in preclinical models and possibly in clinical settings. Deficits in sensory-motor gating, measured by the behavioral paradigm of pre-pulse inhibition, have been documented in all major schizophrenia mouse models, such as DISC1(*68*) and 22q11 chromosome deletion(*28*). Correlative evidence indicates that SG deficits in these mouse models might be related to reduced function or density of inhibitory neurons, in particular of PV-interneurons (e.g.(*28*)). Behavioral sensorimotor gating (paired pulse inhibition) had an important role in preclinical research as the efficacy of neuroleptics in ameliorating this deficit correlates with their clinical efficacy, a property used to screen and identify new molecules such as quetiapine(*13*). However, in absence of knowledge about the specific microcircuit and synaptic mechanism behind SG, this type of research has been largely correlative so far. Our work identifies a synaptic and cellular target responsible for SG at microcircuit scale: *the principal neuron-to-SS-interneuron synapse and its modulation by key molecular players in schizophrenia such as NMDA receptors and dopamine*. The model we propose, summarized in Figure 7, restricts the possible targets behind SG deficits and its clinical consequences. A central role of the dysfunction of SS-interneurons in psychotic disorders is highlighted by recent findings of cell type-specific SS-interneurons abnormalities in postmortem prefrontal tissue of schizophrenic patients(*69*).

Thus, a deficit in the prefrontal-specific developmental increase of NMDA receptors in SS-interneurons, favoring temporal integration at adolescence, might be a key pathophysiological mechanism behind the emergence of schizophrenia prodromic symptoms.

In summary, our data support the view that *a prefrontal deficit of dopaminergic tone impairs NMDA -dependent dendritic integration in pyramidal neuron-SS interneuron synapses and that this could be the key mechanisms behind SG deficits* in psychose-prone disorders such as schizophrenia. The identification of a specific and synaptic cellular target behind a biological endophenotype of psychosis-prone disorders, together with preclinical cell-type specific opto- and chemogenetic tools and spatio-temporally precise brain stimulation clinical techniques, is necessary for new, more targeted and hence possibly more effective treatments of psychotic disorders. Indeed, cell type-specific optogenetics has been now used in the awake primate brain(*70*), and the first optogenetic clinical trials have already been approved in neurology(*71*).

## MATERIALS AND METHODS

### Ethical considerations and animals

Adult male and female C57BL/6J mice (10-13 weeks) were kept at Umeå Center for Comparative Biology with *ad libitum* access to food and water. All procedures were approved by Northern Sweden Ethical Committee (permits A26-2018 and A21-2023). A total of 91 animals were used for this study.

### Surgical procedures

Mice were anaesthetized with 1,5-2,5% isoflurane for virus injections, or sodium pentobarbital (50 mg/kg i.p. induction, 10 mg/kg supplementation) for electrophysiology combined with optogenetics/pharmacology as well as calcium fiberphotometry. For fiberphotometric measurements, a colony of transgenic animals expressing the genetically encoded calcium reporter GCAMP6s was used (strain GP4.3)(*72*). Anesthesia depth was monitored by regularly testing pinch and corneal reflexes, as well breathing rate. The body temperature was kept at 37°C by a rectal probe connected to a heating plate. A local anesthetic (lidocaine) was applied at incision areas. For electrophysiology, a custom-built stereotaxic apparatus was used where a recording chamber was fixed on the skull with dental cement. The craniotomies were performed at following stereotaxic coordinates using bregma as the fixed landmark. OFC: 2,3mm antero-posterior; 1,0mm medio-lateral and 1,6 – 1,9mm dorso-ventral; AAC (−2,8 – - 3,1mm antero-posterior; 4,0mm medio-lateral; 0.6 – 0.8mm dorso-ventral); Frontal Association Cortex (FrA): 2,8mm antero-posterior; 1,0mm medio-lateral; 0,5mm dorso-ventral; and Medial Posterotemporal Asociation Cortex (MPtA, part of the posterior parietal cortex PPC: −2,0mm antero-posterior; 1,0mm medio-lateral, 0,6mm dorso-ventral).

#### Virus injections

The following viruses were injected in OFC (500nl at the rate of 3nl/s, 50nl volume ejection with a delay of 20s between every ejection cycle) using a Nanoject II injector (Drummond Scientific Company, USA) under a BSL-2 stereotaxic hood: AAV1.Ef1a.DIO.ChR2(E123A).EYFP.WPRE.hGH, AAV5-Syn-DIO-stGtACR1-FusionRed and AAV9-CamKII0.4-Cre. After 4 weeks from the injections, optogenetic experiments combined with electrophysiology were performed.

### Histological verifications

Animals were transcardially perfused under deep barbiturate anesthesia (50mg/kg i.p) using 50ml saline followed by 100ml of cold 4% paraformaldehyde (PFA). Post-fixed (1h) brains were transferred to 30% sucrose at 4°C (48h). Brains were coronally sectioned (60μm) on a sliding microtome (Leica Biosystems, Germany), transferred to Superfrost™ Plus slides and coverslipped using Fluoromount-G (Invitrogen, USA). The fluorescent sections were examined with Axio Imager M.2 microscope (Zeiss, Germany). The images were captured at 4x with filter sets 38 (excitation BP 470/40, emission bP525/50 for DiO), filter set 49 (excitation G365, emission BP 445/50 for DAPI) and filter set 43HE (excitation 550/25 HE, emission 605/70 HE for DiI). All frames were stitched together using the Zeiss Zen software. Franklin and Paxinos atlas(*73*) was used to identify brain/cortical areas.

### *In-vivo* electrophysiological recordings

Borosilicate glass pipettes (1,5mm outer diameter) filled with saline (0.9% NaCl) or with the drug of interest were inserted in a 0.5 x 0.5mm craniotomy in OFC or AAC. For the experiments in which pharmacology and /or optogenetics were combined the pipette tip was broken to a tip diameter of 12-15μm, corresponding to a resistance of 0,1MΩ. Sound stimuli were presented 50 times and acoustic evoked potentials were recorded using an EPC10 amplifier (HEKA, Germany) in current clamp after band-pass-filtering (0.1–100Hz) and amplification (1000X) (NPI Electronik, Germany). The signal was digitized at 1kHz and recorded via Patchmaster software (HEKA, Germany).

#### LFP signal analysis

averaged signals were baseline-corrected (200ms pre stimulus response window). The peak amplitude in the OFC was searched within a 400ms window, 200ms after stimulus onset. For AAC, peak amplitudes were taken within an equally long 400ms time window starting immediately after stimulus onset instead. In some cases (25 out of 32, a small photoelectric voltage drop was subtracted offline from the averaged evoked potential trace from light onset as an abrupt, light-locked voltage drop occurring within the first few ms after light (median: 3,0ms; see example in Supplementary Figure 3B).

### Pharmacology

The following drugs were used: 0,6 μl of Gabazine (1,5μM) and/ or CGP 52432 (1μM) (Tocris); 0,6μl of Flupentixol dihydrochloride (5mM, Merck) and 0,6μl of D-AP5 (2,5mM; Tocris). Glass micropipettes (15μm outer tip) filled with drugs were inserted under negative pressure to avoid leakage of the drug before injection. Once a pre-drug baseline recording was obtained, the drug was carefully delivered under visual inspection of the meniscus by delivering 100-200mm Hg pressure pulses over 5min using a controlled pressure-injection module (NIM, Heidelberg, Germany). Post-drug local field potentials (no pipette movement) were recorded 30min after each drug injection.

### Optogenetics

For optogenetic experiments, combined with electrophysiology, we used a LED light source (Prizmatix), coupled with a 200μm, 0,22NA optical fiber (Thor Labs) that was inserted via a dedicated 30° angled port in the electrode pipette holder (Optopatcher, ALA Scientific). The optic fiber tip was positioned to the lowermost possible position in the recording pipette (12-15μm outer tip diameter), at an approximate distance from the recording pipette tip of 150-200μm. A power of 0,8 - 0,9mW was used for all experiments (power density: 6-7mW/mm^2^) and the wavelengths used were 520nm for the experiments with the GTCAR1 opsin and 460nm for the experiments with the ChR2 opsin.

### Calcium fiberphotometry

Fiberphotometric measurements *in-vivo* were performed by slowly advancing the bare optic fiber tip in OFC at 50% intensity compared to the one used during the actual measurements. The fiber was placed at about 1,2-1,5mm below the brain surface. Once in place, we waited for 2-3min with the light off before increasing the intensity to 100% (that was done just before starting the recording). The fiberphotometric device was a commercial system from NPI elektronik (FOM-02M-Fiber OptoMeter II) containing LEDs for excitation and a photomultiplier tube (PMT). Excitation was provided by a blue LED (wavelength 450-490nm), connected to an optic fiber (200µm, 0,39NA, Thor Labs), used at a power setting so to provide 0,9mW at the fiber output (corresponding to a power density of 7mW/mm^2^). The emission fluorescence light was collected in the 500-540nm band (pre PMT band-pass filter). The PMT voltage output was band-passed filtered between 0.1-50Hz and amplified 100 times, digitized at 1KHz and acquired on the Patchmaster software. When calcium photometry was combined with pharmacological injections of a drug, a second manual 3D micromanipulator (Narishige M-3333) with the drug filled glass pipette was positioned in a second craniotomy (3mm from midline, same antero-posterior level as the recording craniotomy) so as to drug-perfuse the region of interest (the estimated distance between the optic fiber tip and the injecting pipette being indeed ca 100 – 200µm).

#### Calcium-related PMT signal

The calcium-related PMT voltage signal was averaged for each experiment and the peak signals (minima) were measured between 200 and 600ms from sound stimulus onset on each experimental condition average trace (50 sweeps were averaged for calcium fiberphotometry to measure sound-evoked responses, 100 sweeps to measure SG). All traces were processed post-acquisition using the adjacent average smoothening with 50 points of window before the peak amplitudes were measured.

### Auditory stimulation

Pure tones were generated by Matlab, range 4–64kH (4, 8, 11, 16, 22, 32, 44, 50, 64KHz), 75dB, duration: 400ms, 5s interstimulus interval. Tones were analogically converted using an RX6 MultiFunction Processor and delivered randomly through TDT ED1 speakers (Tucker-Davis Technologies, USA), located 10cm from the contralateral ear. Auditory evoked potentials were performed before the SG test to ensure the presence of reliable and solid acoustic response. The threshold for ultrasound definition was 20kHz.

#### Sensory Gating protocol

A paired stimulus tested for SG in both OFC and the AAC. First, the optimal frequency between the range 4-64kHz, 75dB was identified. The protocol was written in RPvdsEx software of RX6 MultiFunction Processor (TDT technologies, US) and followed sensory gating paradigm in rodent medial prefrontal cortex(*18*). Each repetition consisted of a 200ms prestimulus followed by two identical 100ms stimulus pulses (doublet), which were separated by a 500ms interpulse interval and finally a 700ms poststimulus block. There was an intertrial interval of 10s. Responses were averaged over 100 repetitions and baseline corrected (200ms of pre stimulus time window). For determining the optimum interpulse interval, animals (N=4) were also tested at interpulse intervals of 200, 500, 800 and 1100ms respectively, and in this latter case only 50 repetitions per interstimulus time were measured.

##### Computational modeling of optogenetic activation and inhibition experiments

A phenomenological model based on population firing rate (*R_k_*) equations (I) was constructed, and a 5-node network was simulated, corresponding to five distinct interacting neuron populations with corresponding distinct time constants *τ*_*k*_: auditory cortex cells (AAC), pyramidal cells at the orbitofrontal cortex (PYR), somatostatin-positive interneurons at the orbitofrontal cortex (SST), parvalbumin-positive interneurons at the orbitofrontal cortex (PV), and vasointestinal peptide-positive interneurons at the orbitofrontal cortex (VIP).

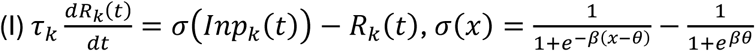

The sum of time-dependent inputs *Inp*_*k*_(*t*) (II) at the AAC and PYR nodes were postulated to correspond to experimental LFPs measured at the auditory cortex and orbitofrontal cortex, respectively. Parameters *A_k_* were coefficients modulating the effect of optogenetic manipulation in each node, and *W_kj_* represented connection strengths between pairs of nodes.

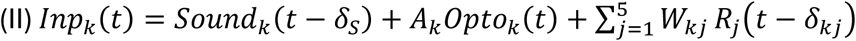

A time delay of *δ*_1,0_ = *δ*_3,0_ = 103*ms* was imposed over the AAC-to-PYR and AAC-to-PV connections, in order to match experimental observations of deflection onset times. All other connections involved no time delays (*δ*_(*k*,*j*)∉{(1,0),(3,0)}_ = 0*ms*).

Sound stimuli were simulated as short square-shaped pulses, activated at times *t*_1_ and *t*_2_ with a duration of *Δ*_*s*_ = 20*ms* and intensity of 0.1 arbitrary units (III), acting exclusively on the AAC population. A fixed delay of *δ*_*S*_ = 12*ms* was imposed between the sound stimulation times and the corresponding onsets of *Sound*_0_(*t*) at the AAC node, to match experimental onset times. No other node received direct inputs corresponding to the sound stimulation (*Sound*_*k*≠0_(*t*) = 0).

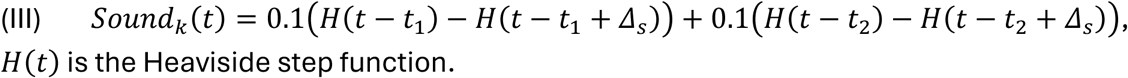

Optogenetic manipulations were simulated as additional input terms acting on the corresponding populations, with either positive or negative sign for optogenetic activation or inhibition, respectively. The onset time for optogenetic stimulation matched the time *t*_1_of the first sound pulse (with no transmission delay), and the intensity peaked at 0.1 arbitrary units but was allowed to decay after onset according to the functional form in (IV), with a characteristic time of of *δ*_*op*_ = 10*ms*, down to a level *B*_*ACT*_ or *B*_*INH*_ that was different for activation or inhibition experiments to partly account for possible differences between the different opsins.

For each different experimental condition, optogenetic inputs were delivered only to the node corresponding to the target population (either SST [*k* = 2] or PV [*k* = 3] with *Opto*_*k*_(*t*) = 0 for all other *k*), and with the appropriate signs and asymptotes *B*_*ACT*_ or *B*_*INH*_ corresponding to either activation or inhibition.

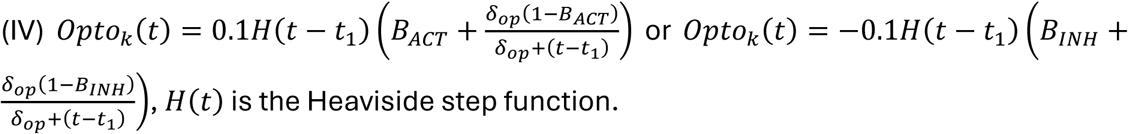

Time series were integrated with the Euler method with a 0.1*ms* time step. Experimental LFPs were upsampled to a matching 0.1*ms* sampling rate, and OFC LFPS were collectively normalized such that the baseline LFP peaked at an arbitrary value of 0.1, and all other conditions maintained the corresponding relative intensity with respect to the baseline. The AAC LFP was independently normalized to peak at the same arbitrary value of 0.1.

Free parameters (*A*_*k*_, *B*_*ACT*_, *B*_*INH*_, *τ*_*k*_, *W*_*kj*_, 25 in total) were found via the simultaneous optimization over all five different experimental conditions of similarity functions based on the either the local maximization of the Pearson correlation or local minimization of the sum of squared differences between the simulated time-dependent inputs *Inp*_*k*_(*t*) and the observed LFPs at the OFC and AAC (but note that AAC LFPs were only available for the no-stimulation condition).

The selected model instance, which displayed the best combination of reproducing LFP shapes and gating ratios for all conditions simultaneously, was used to perform a blind inference of the dependence between baseline gating ratio and inter-stimulus interval, a separate experimental result that was not included in the model training. The observed correspondence between model inference and experimental result was taken as confirmation of the biological plausibility of the model results.

### Statistical analysis of physiological data

Normality was tested by Kolmogorov Smirnov tests. Paired T-tests compared paired data (mean ± SEM are shown), whereas unpaired data comparisons were done by t-test or by ANOVA followed by Bonferronís posthoc tests. Significance was at 0.05 level. Statistical analyses were done using OriginPro 2017 (Origin Lab, USA).

## ABBREVIATIONS

SG: sensory gating
LFP: local field potential
OFC: orbitofrontal cortex
SS: somatostatin
PV: parvalbumin
PMT: photomultiplier
AAC: acoustic association cortex

---

**Figure S1.**
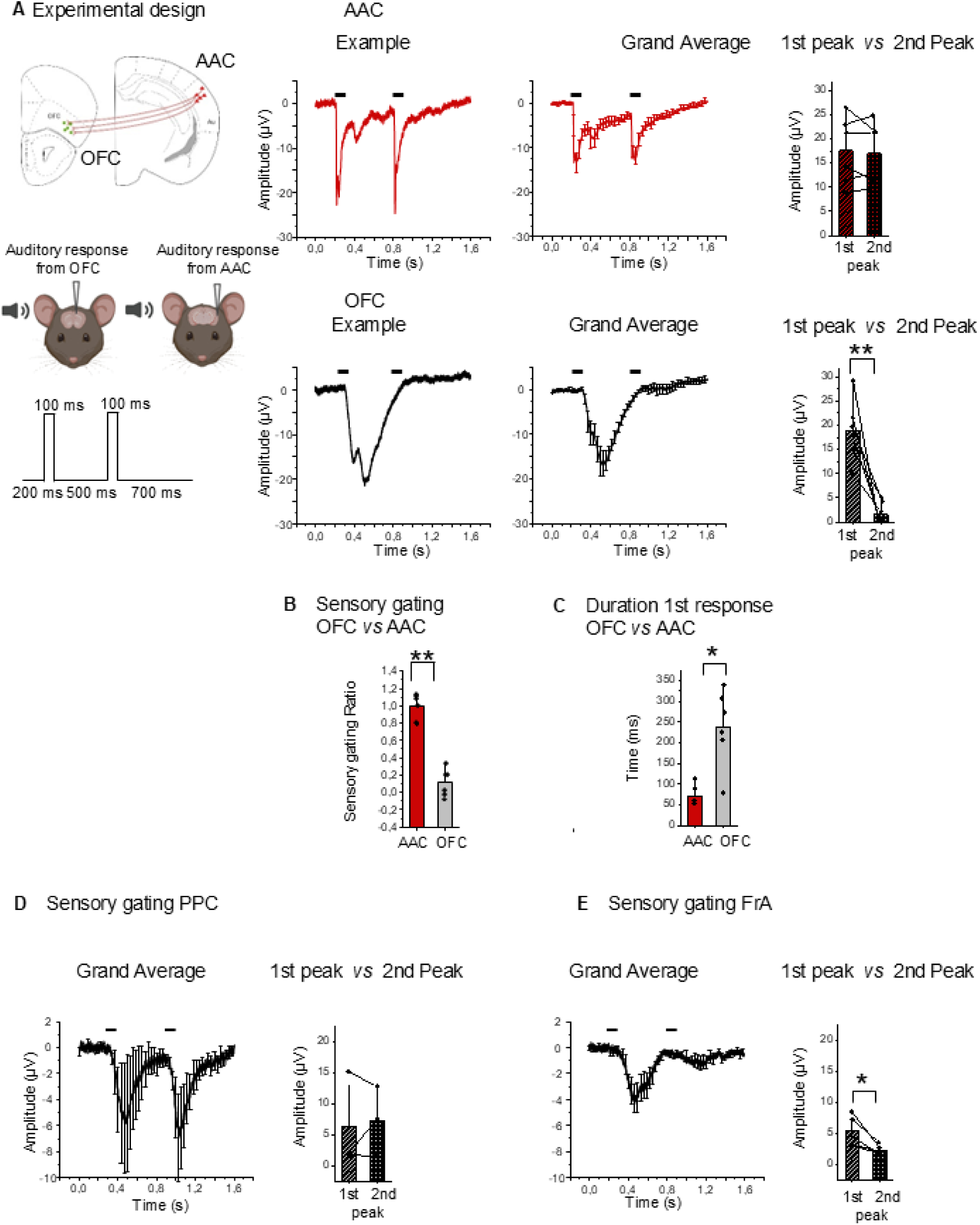
SG is dramatic in mouse OFC and prefrontal cortex but negligible in its input acoustic cortex and other associative cortical areas. **A**. AAC-to-OFC connectivity and experiment sketch of the dual recording in AAC and OFC. Examples (left plot) and grand averages (middle plot) of SG measurements in AAC (top row) and OFC (bottom). Paired quantifications of the first and second peak amplitudes in the right plots for AAC (1^st^ peak: 17,5 ± 2,9μV vs 2^nd^ peak: 16,9 ± 2,6μV; paired t-tests, p=0,63) and OFC (1^st^ peak: 18,9 ± 2,6μV vs 2^nd^ peak: 1,6 ± 1,1μV; paired t-test **p=0,004), respectively). **B**. SG ratio values measured in OFC were statistically smaller compared to AAC (OFC: 0,12 ± 0,07 vs AAC: 0,99 ± 0,06; paired t-test, **p=0,001). **C**. The duration of the first sound response (expressed as duration at half peak amplitude) was larger in OFC compared to AAC (OFC: 238,5 ± 37,9ms vs AAC: 70,0 ± 10,3ms; t-test, *p=0,013). **D,E**. Grand averages and SG ratios upon SG protocol run in posterior parietal cortex (PPC, left plot and histograms-1^st^ peak vs 2^nd^ peak: 6,31± 4,4μV vs 7,23 ± 3,25μV) and in the frontal association area (FrA, right plot and histograms-1^st^ peak vs 2^nd^ peak: 5,27 ± 1,09μV vs 2,27 ± 0,37μV). No significant SG was measured in PPC (paired t-test; p=0,733), whereas there was a robust SG in FrA (paired t-test; *p=0,025).

**Figure S2.**
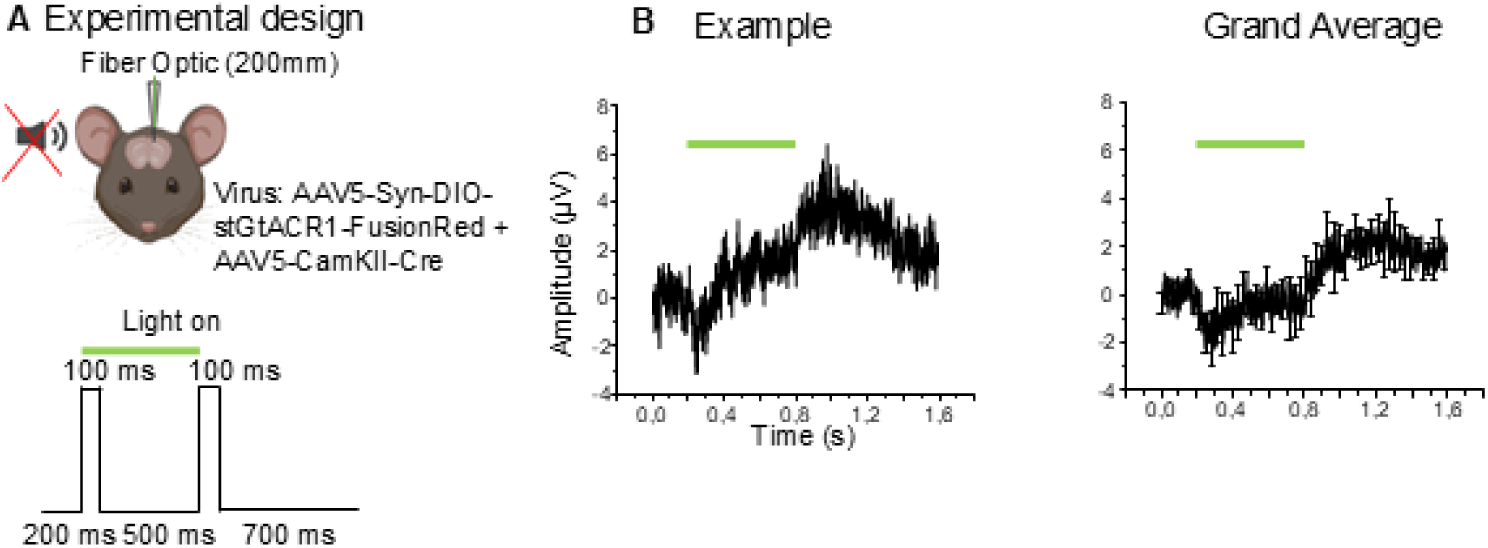
Control experiment showing absence of rebound downward, excitatory LFP signal upon switching off optogenetic stimulation in the context of experiment of main Figure 2A-C. **A**. Experiment sketch. **B**. Example and grand average of the data.

**Figure S3.**
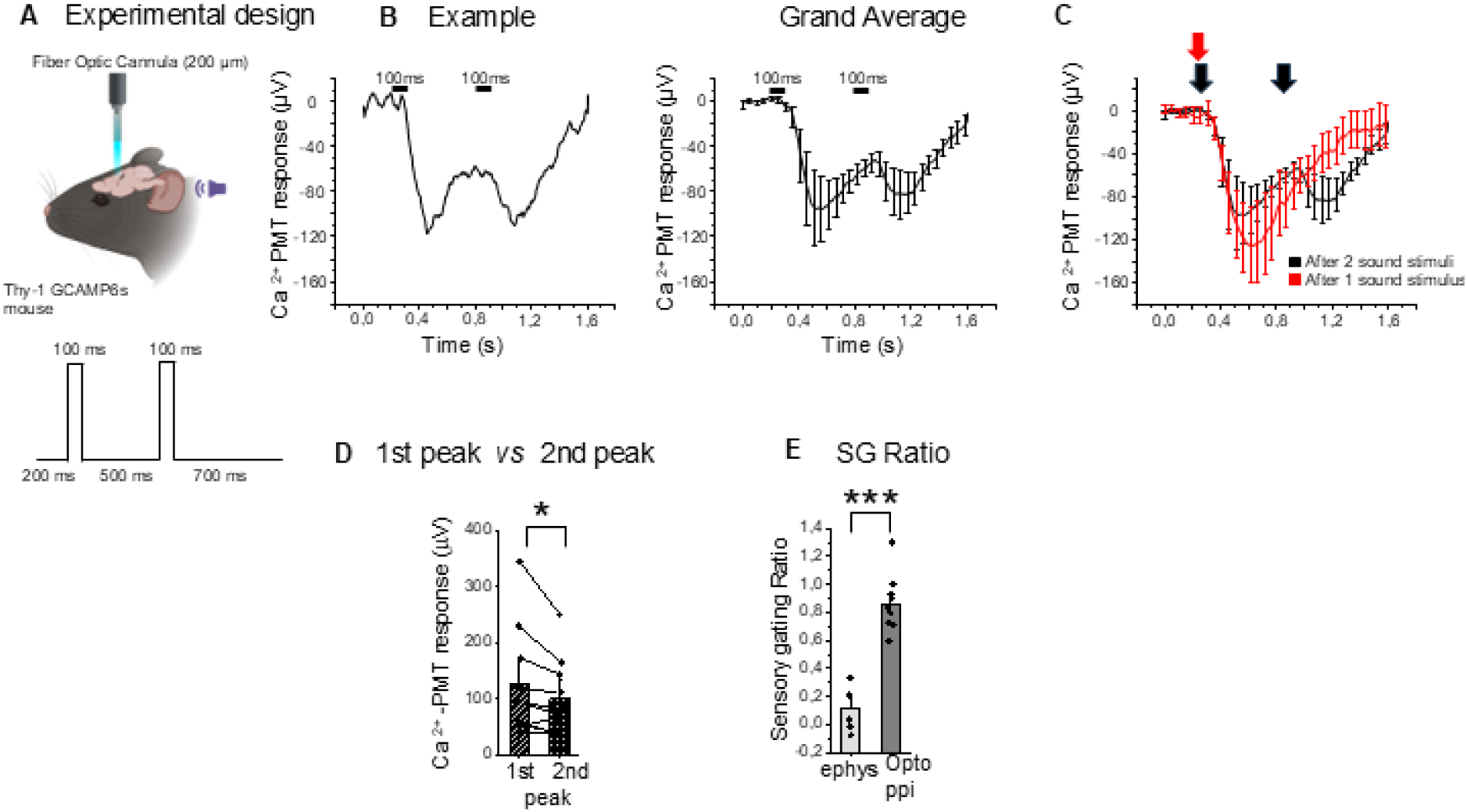
SG measured by neuropil calcium fiberphotometry in Thy1-GCAMP6s mice is smaller compared to the one measured with electrophysiology. **A.** Experiment sketch: an optic fiber was inserted in OFC in mice expressing the genetically encoded calcium indicator GCAMP6s in Thy1 positive excitatory neurons of OFC during anesthesia. **B.** Calcium-related, voltage PMT signal recorded during one example of SG protocol (example on the left plot, grand-average on the right plot). **C.** Comparison of the time course of the calcium signal in response to SG protocol and to a single acoustic pulse. Note that the calcium-related signal does not reach the baseline upon 1^st^ sound stimulus. **D.** SG is detectable with calcium fiberphotometry but is significantly smaller than the one recorded with electrophysiology. Left: the 2^nd^ peak is significantly smaller than the 1^st^ (1^st^ response-124,9 ± 30,9µV vs 2^nd^ response-100,2 ± 21,7µV; paired t-test, *p=0,04). Right: SG ratios measured with electrophysiology (ephys) are significantly smaller than the ones measured optically with calcium fiberphotometry (opto; SG ratio ephys: 0,12 ± 0,07 *vs* SG ratio opto: 0,85 ± 0,06; t-test, ***p=0,000002).

**Figure S4. LFP signal driven by SS-interneurons stimulation per se in OFC and effects of extracellular GABA blockade. A**. Grand average of experiments in which the green LED source and the optic fiber were inserted in SS-Cre mice and switched on in absence of any acoustic stimulus (N=3). These data show that the initial upward deflection is related to opsin activation *per se.* The red trace is the optically-evoked LFP looked after injection of extracellular GABA blockers (Gabazine 3μM and CGP532 1μM, 600nL). Note the decrease of the first upward peak. **B**. Correction of photoelectric artifact. The tiny voltage drop caused by the light onset (see right inset) was subtracted from the recorded sweep.

**Figure S5. Photoinhibition of SS-interneurons during the first time window has effects on the second response similar to photoinhibition during the entire protocol. A.** Experiment sketch. Comparison of SG upon photoinhibition of PV-(green) or SS-(blue) interneurons when the light was one during the first time window (black lines in plots) or during both time window (colored lines in plots). **B, C.** Comparison of grand averages (**B**) and quantifications of 1^st^, 2^nd^ responses and SG ratios (**C**) between the two cases. None of the comparisons were significant. Note also the similarity of the patterns in case of photoinhibition of SS interneurons (bottom row, blue vs black lines for illumination during the whole sweep or during the first time window, respectively). Similar results are true for the PV-interneurons.

**Figure S6.**
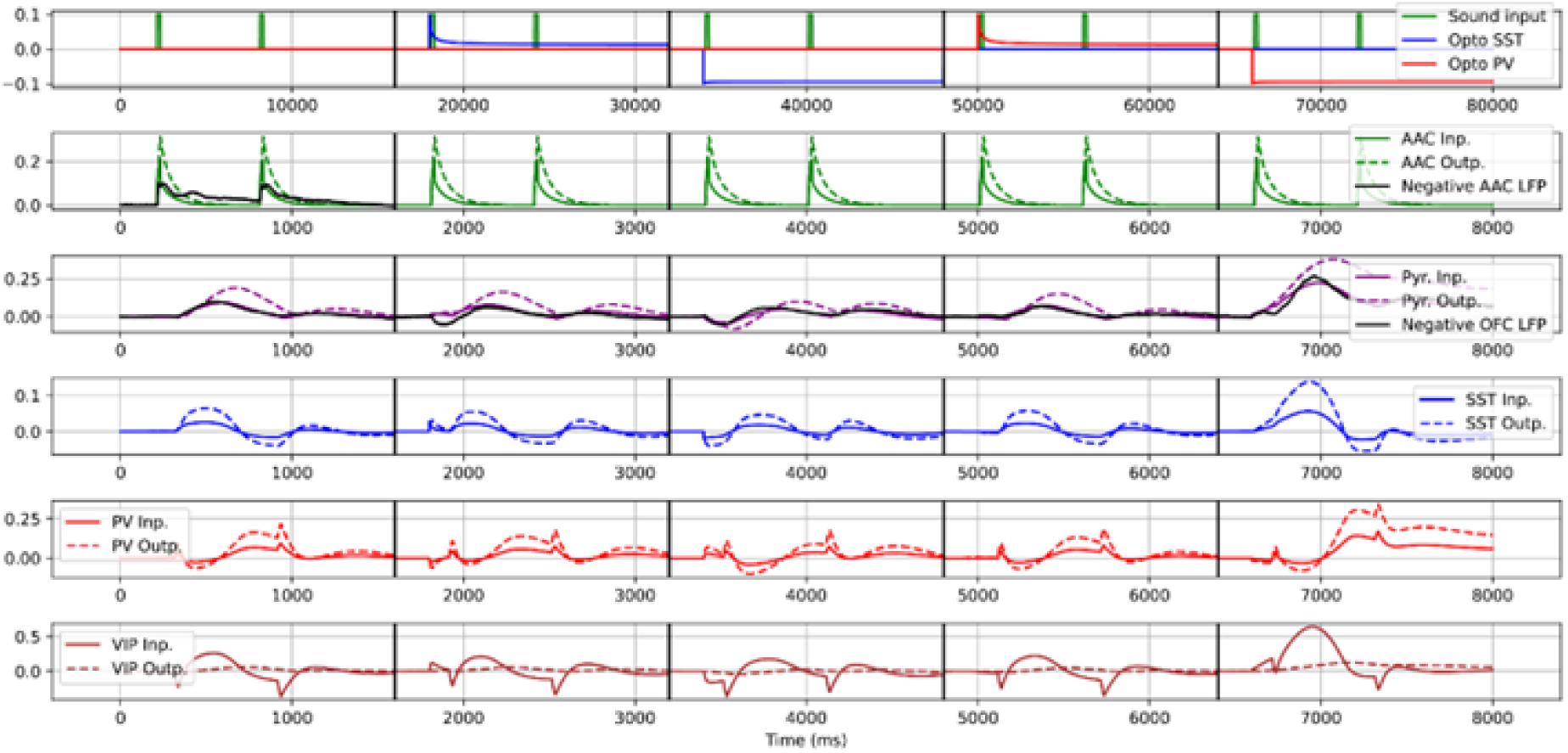
Time-dependent inputs and outputs for all simulated nodes at the five different experimental conditions. First row: magnitude of sound inputs and optogenetic manipulations. Subsequent rows: magnitude of total inputs (continuous lines) and outputs (dashed lines) at the AAC, PYR, SST and VIP nodes, respectively. Where present, black lines correspond to normalized experimental curves. Conditions (left to right): baseline, optogenetic excitation of SST cells, optogenetic inhibition of SST cells, optogenetic excitation of PV cells, optogenetic inhibition of PV cells.

**Figure S7.**
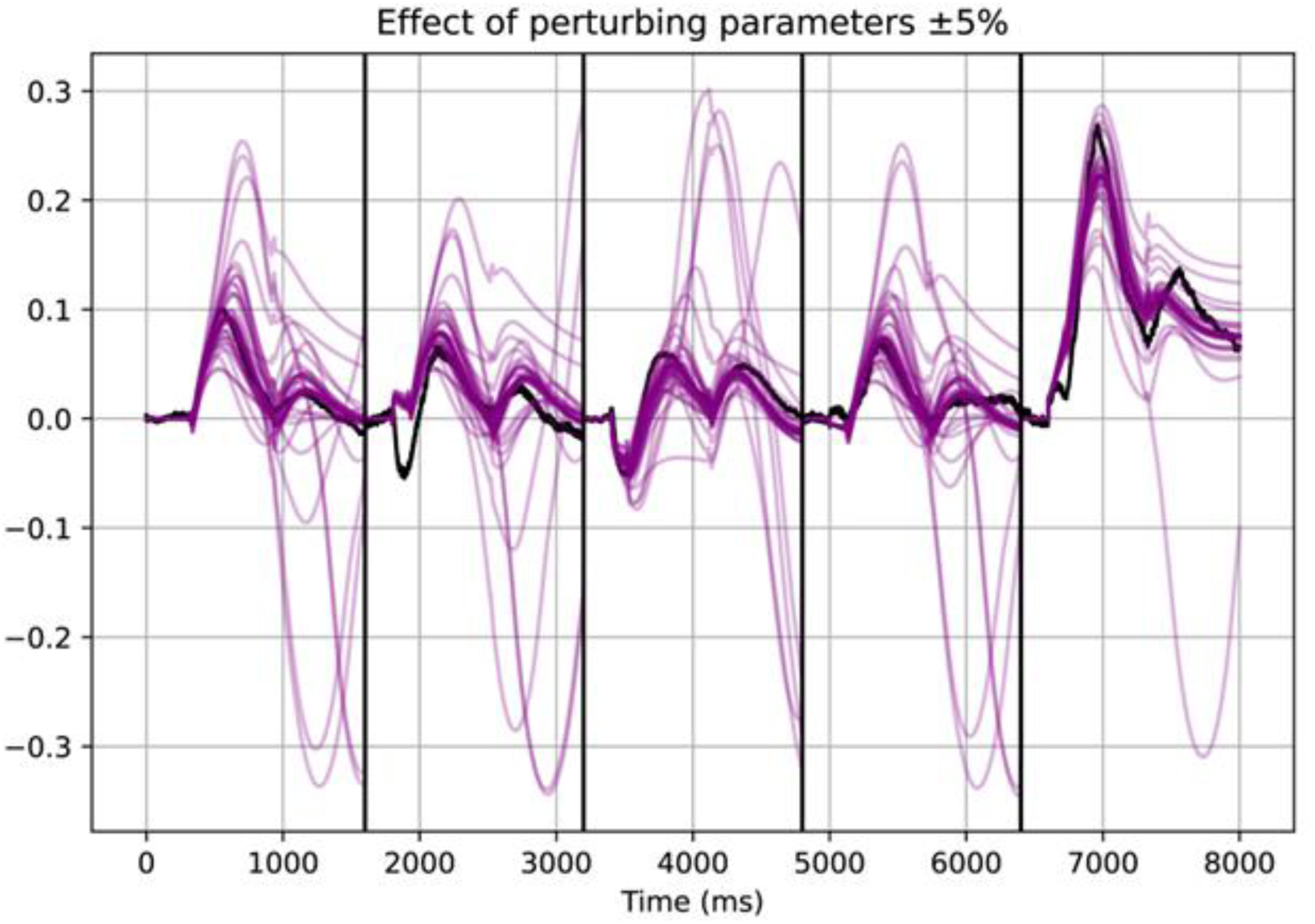
Time-dependent inputs at the PYR node corresponding to each individual run with a different perturbed parameter. Concatenated conditions, left to right: baseline, optogenetic excitation of SST cells, optogenetic inhibition of SST cells, optogenetic excitation of PV cells, optogenetic inhibition of PV cells.

**Figure S8.**
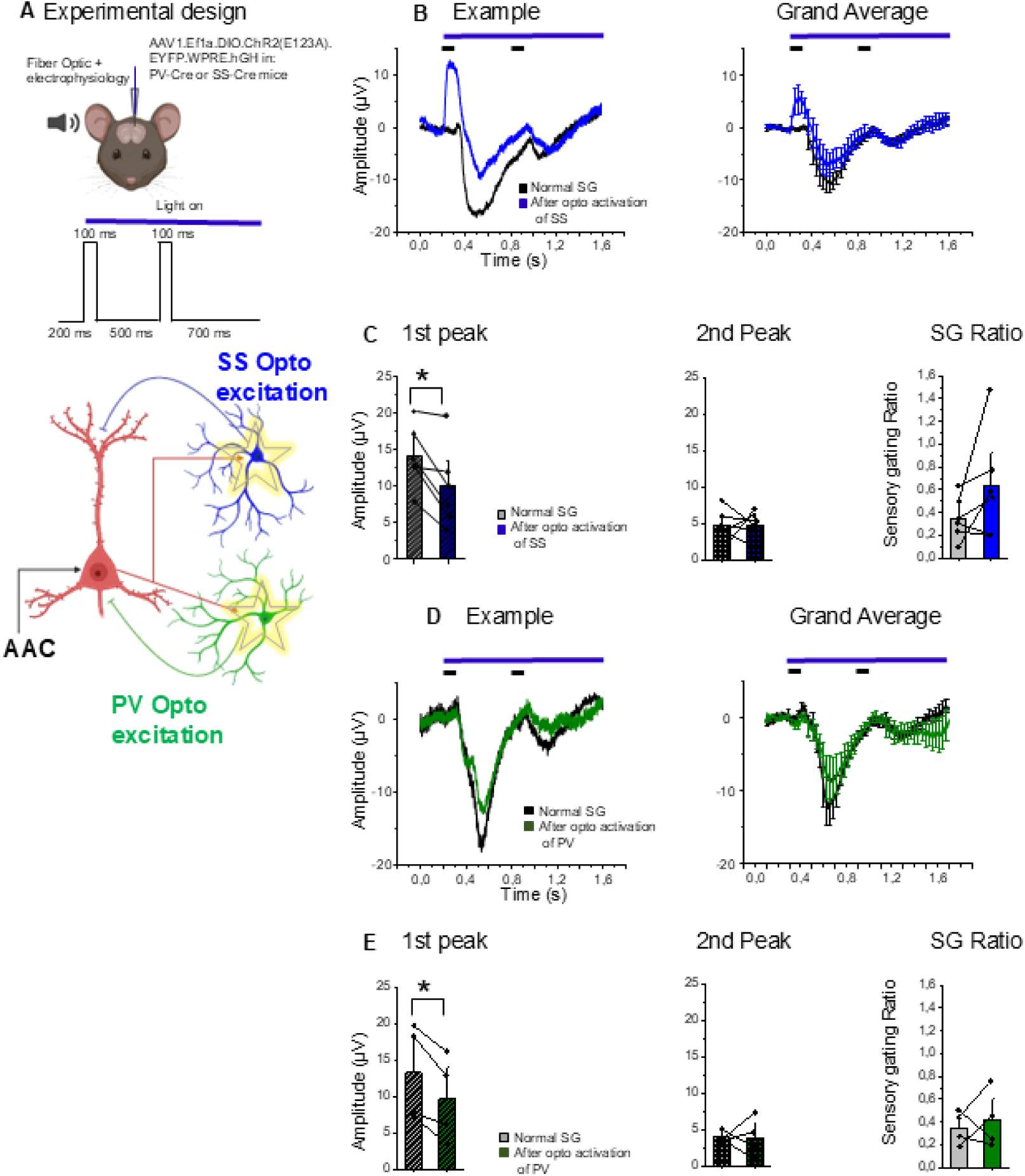
Documentation of the effects of photostimulation of GABAergic interneurons *per se* on SG. **A.** Experiment sketch: PV- or SS-interneurons were optogenetically stimulated by selective, Cre-dependent expression of ChR2 in corresponding transgenic mice. **B.** Examples and grand averages of LFP responses during SG upon photostimulation of SS-interneurons (black vs blue for responses without vs with photostimulation, respectively). Note the reduction of the first sound response without any increase in the 2^nd^ peak response. **C.** Quantification of the results. Note the significant reduction of the 1^st^ sound response (N=6; 1^st^ peak-before *vs* after opto activation: 12,8 ± 1,8µV *vs* 9,9 ± 2,4µV; paired t-test, *p=0,02; 2^nd^ peak-before vs after optoactivation: 4,5 ± 0,8µV *vs* 4,7 ± 0,8µV; p =0,96; SG ratio-before *vs* after optoactivation: 0,38 ± 0,07 *vs* 0,63 ± 0,19; paired t-test, p=0,15). **D.** Examples and grand averages of LFP responses during SG upon photostimulation of PV-interneurons (black vs green for responses without vs with photostimulation, respectively). Note the reduction of the 1st sound response. **E.** Quantification of the results. Note the significant reduction of the 1^st^ sound response (1^st^ peak: before *vs* after optoactivation: 13,2 ± 3,3µV *vs* 9,6 ± 2,9µV; paired t-test, *p=0,01; 2^nd^ peak: before and after photoactivation of PV: 3,9 ± 0,3µV *vs* 3,8 ± 1,4µV; paired t-test, p=0,96; SG ratio-before and after photoactivation: 0,35 ± 0,07 *vs* 0,42 ± 0,13; paired t-test, p=0,66).

## AUTHOR CONTRIBUTIONS

A.T. performed and analyzed all the experiments and contributed critically to project development; T.S. contributed to the parvalbumine optogenetic experiments and performed the histology, L.C. designed and performed all the modelling work, O.L. gave intellectual contributions to the manuscript, P.M. conceived and supervised the project and wrote the manuscripts with critical contributions from all authors.

## ACKNOWLEDGMENTS

we are grateful to Per Utsi, Roger Widmark, Anders Bäckström and Göran Dahlgren for excellent technical assistance, and to Staffan Johansson and Per Petersson for critically reading the manuscript.

## Notes

### Competing Interest Statement

The authors have declared no competing interest.

## REFERENCES

1. H. Kim, S. Ahrlund-Richter, X. Wang, K. Deisseroth, M. Carlen, Prefrontal Parvalbumin Neurons in Control of Attention. Cell 164, 208–218 (2016).

2. J. A. Micoulaud-Franchi et al., Sensory gating in adult with attention-deficit/hyperactivity disorder: Event-evoked potential and perceptual experience reports comparisons with schizophrenia. Biol Psychol 107, 16–23 (2015).

3. P. Lichtenstein et al., Common genetic determinants of schizophrenia and bipolar disorder in Swedish families: a population-based study. Lancet 373, 234–239 (2009).

4. D. Bipolar, d. r. v. e. Schizophrenia Working Group of the Psychiatric Genomics Consortium. Electronic address, D. Bipolar, C. Schizophrenia Working Group of the Psychiatric Genomics, Genomic Dissection of Bipolar Disorder and Schizophrenia, Including 28 Subphenotypes. Cell 173, 1705–1715 e1716 (2018).

5. L. E. Adler et al., Neurophysiological evidence for a defect in neuronal mechanisms involved in sensory gating in schizophrenia. Biol Psychiatry 17, 639–654 (1982).

6. C. Siegel, M. Waldo, G. Mizner, L. E. Adler, R. Freedman, Deficits in sensory gating in schizophrenic patients and their relatives. Evidence obtained with auditory evoked responses. Arch Gen Psychiatry 41, 607–612 (1984).

7. E. M. Sanchez-Morla et al., P50 sensory gating deficit is a common marker of vulnerability to bipolar disorder and schizophrenia. Acta Psychiatr Scand 117, 313–318 (2008).

8. H. K. Hamilton et al., Clinical and Cognitive Significance of Auditory Sensory Processing Deficits in Schizophrenia. Am J Psychiatry 175, 275–283 (2018).

9. C. L. Shen et al., P50, N100, and P200 Auditory Sensory Gating Deficits in Schizophrenia Patients. Front Psychiatry 11, 868 (2020).

10. L. A. Jones, P. J. Hills, K. M. Dick, S. P. Jones, P. Bright, Cognitive mechanisms associated with auditory sensory gating. Brain Cogn 102, 33–45 (2016).

11. A. Khani et al., Large-Scale Networks for Auditory Sensory Gating in the Awake Mouse. eNeuro 6, (2019).

12. J. Smucny, K. E. Stevens, A. Olincy, J. R. Tregellas, Translational utility of rodent hippocampal auditory gating in schizophrenia research: a review and evaluation. Transl Psychiatry 5, e587 (2015).

13. N. R. Swerdlow, G. A. Light, Animal Models of Deficient Sensorimotor Gating in Schizophrenia: Are They Still Relevant? Curr Top Behav Neurosci 28, 305–325 (2016).

14. L. E. Adler, G. Rose, R. Freedman, Neurophysiological studies of sensory gating in rats: effects of amphetamine, phencyclidine, and haloperidol. Biol Psychiatry 21, 787–798 (1986).

15. F. Koukouli et al., Nicotine reverses hypofrontality in animal models of addiction and schizophrenia. Nat Med 23, 347–354 (2017).

16. K. M. Hershman, R. Freedman, P. C. Bickford, GABAB antagonists diminish the inhibitory gating of auditory response in the rat hippocampus. Neurosci Lett 190, 133-136 (1995).

17. A. Tripathi, S. S. Sato, P. Medini, Cortico-cortical connectivity behind acoustic information transfer to mouse orbitofrontal cortex is sensitive to neuromodulation and displays local sensory gating: relevance in disorders with auditory hallucinations? J Psychiatry Neurosci 46, E371–E387 (2021).

18. R. P. Mears, A. C. Klein, H. C. Cromwell, Auditory inhibitory gating in medial prefrontal cortex: Single unit and local field potential analysis. Neuroscience 141, 47–65 (2006).

19. G. M. Olsen et al., Organization of Posterior Parietal-Frontal Connections in the Rat. Front Syst Neurosci 13, 38 (2019).

20. M. Mahn et al., High-efficiency optogenetic silencing with soma-targeted anion-conducting channelrhodopsins. Nat Commun 9, 4125 (2018).

21. C. L. Miller et al., Phencyclidine and auditory sensory gating in the hippocampus of the rat. Neuropharmacology 31, 1041–1048 (1992).

22. T. Fellin et al., Endogenous nonneuronal modulators of synaptic transmission control cortical slow oscillations in vivo. Proc Natl Acad Sci U S A 106, 15037–15042 (2009).

23. S. L. Smith, I. T. Smith, T. Branco, M. Hausser, Dendritic spikes enhance stimulus selectivity in cortical neurons in vivo. Nature 503, 115–120 (2013).

24. H. Dana et al., Thy1-GCaMP6 transgenic mice for neuronal population imaging in vivo. PLoS One 9, e108697 (2014).

25. C. Montinaro et al., Influence of the anatomical features of different brain regions on the spatial localization of fiber photometry signals. Biomed Opt Express 12, 6081–6094 (2021).

26. H. Jeong, D. Kim, M. Song, S. B. Paik, M. W. Jung, Distinct roles of parvalbumin- and somatostatin-expressing neurons in flexible representation of task variables in the prefrontal cortex. Prog Neurobiol 187, 101773 (2020).

27. D. Kim et al., Distinct Roles of Parvalbumin- and Somatostatin-Expressing Interneurons in Working Memory. Neuron 92, 902–915 (2016).

28. A. Tripathi et al., Cognition- and circuit-based dysfunction in a mouse model of 22q11.2 microdeletion syndrome: effects of stress. Transl Psychiatry 10, 41 (2020).

29. S. M. Perez, A. Boley, D. J. Lodge, Region specific knockdown of Parvalbumin or Somatostatin produces neuronal and behavioral deficits consistent with those observed in schizophrenia. Transl Psychiatry 9, 264 (2019).

30. G. Iurilli et al., Sound-driven synaptic inhibition in primary visual cortex. Neuron 73, 814–828 (2012).

31. N. Wagatsuma, S. Nobukawa, T. Fukai, A microcircuit model involving parvalbumin, somatostatin, and vasoactive intestinal polypeptide inhibitory interneurons for the modulation of neuronal oscillation during visual processing. Cereb Cortex 33, 4459–4477 (2023).

32. A. J. Keller et al., A Disinhibitory Circuit for Contextual Modulation in Primary Visual Cortex. Neuron 108, 1181–1193 e1188 (2020).

33. G. Buzsaki, C. A. Anastassiou, C. Koch, The origin of extracellular fields and currents--EEG, ECoG, LFP and spikes. Nat Rev Neurosci 13, 407–420 (2012).

34. M. J. Dunn, S. Killcross, Medial prefrontal cortex infusion of alpha-flupenthixol attenuates systemic d-amphetamine-induced disruption of conditional discrimination performance in rats. Psychopharmacology (Berl) 192, 347-355 (2007).

35. K. U. S. Islam, N. Meli, S. Blaess, The Development of the Mesoprefrontal Dopaminergic System in Health and Disease. Front Neural Circuits 15, 746582 (2021).

36. T. Ott, A. Nieder, Dopamine and Cognitive Control in Prefrontal Cortex. Trends Cogn Sci 23, 213–234 (2019).

37. F. Luo, A. Sclip, S. Merrill, T. C. Sudhof, Neurexins regulate presynaptic GABA(B)-receptors at central synapses. Nat Commun 12, 2380 (2021).

38. N. R. Wilson, C. A. Runyan, F. L. Wang, M. Sur, Division and subtraction by distinct cortical inhibitory networks in vivo. Nature 488, 343–348 (2012).

39. J. F. Sturgill, J. S. Isaacson, Somatostatin cells regulate sensory response fidelity via subtractive inhibition in olfactory cortex. Nat Neurosci 18, 531–535 (2015).

40. S. El-Boustani, M. Sur, Response-dependent dynamics of cell-specific inhibition in cortical networks in vivo. Nat Commun 5, 5689 (2014).

41. M. Krause, W. E. Hoffmann, M. Hajos, Auditory sensory gating in hippocampus and reticular thalamic neurons in anesthetized rats. Biol Psychiatry 53, 244–253 (2003).

42. R. Cools, M. D’Esposito, Inverted-U-shaped dopamine actions on human working memory and cognitive control. Biol Psychiatry 69, e113–125 (2011).

43. D. Di Domenico, L. Mapelli, Dopaminergic Modulation of Prefrontal Cortex Inhibition. Biomedicines 11, (2023).

44. W. J. Gao, P. S. Goldman-Rakic, Selective modulation of excitatory and inhibitory microcircuits by dopamine. Proc Natl Acad Sci U S A 100, 2836–2841 (2003).

45. J. Cousineau et al., Dopamine D2-Like Receptors Modulate Intrinsic Properties and Synaptic Transmission of Parvalbumin Interneurons in the Mouse Primary Motor Cortex. eNeuro 7, (2020).

46. K. Chen, G. Yang, K. F. So, L. Zhang, Activation of Cortical Somatostatin Interneurons Rescues Synapse Loss and Motor Deficits after Acute MPTP Infusion. iScience 17, 230–241 (2019).

47. J. M. Fuster, G. E. Alexander, Neuron activity related to short-term memory. Science 173, 652–654 (1971).

48. K. Kubota, H. Niki, Prefrontal cortical unit activity and delayed alternation performance in monkeys. J Neurophysiol 34, 337–347 (1971).

49. X. J. Wang, 50 years of mnemonic persistent activity: quo vadis? Trends Neurosci 44, 888–902 (2021).

50. M. Wang et al., NMDA receptors subserve persistent neuronal firing during working memory in dorsolateral prefrontal cortex. Neuron 77, 736–749 (2013).

51. S. Vijayraghavan, M. Wang, S. G. Birnbaum, G. V. Williams, A. F. Arnsten, Inverted-U dopamine D1 receptor actions on prefrontal neurons engaged in working memory. Nat Neurosci 10, 376–384 (2007).

52. M. E. Larkum, T. Nevian, M. Sandler, A. Polsky, J. Schiller, Synaptic integration in tuft dendrites of layer 5 pyramidal neurons: a new unifying principle. Science 325, 756–760 (2009).

53. L. Goetz, A. Roth, M. Hausser, Active dendrites enable strong but sparse inputs to determine orientation selectivity. Proc Natl Acad Sci U S A 118, (2021).

54. A. Reyes et al., Target-cell-specific facilitation and depression in neocortical circuits. Nat Neurosci 1, 279–285 (1998).

55. A. Pala, C. C. H. Petersen, In vivo measurement of cell-type-specific synaptic connectivity and synaptic transmission in layer 2/3 mouse barrel cortex. Neuron 85, 68–75 (2015).

56. M. H. Kim et al., Target cell-specific synaptic dynamics of excitatory to inhibitory neuron connections in supragranular layers of human neocortex. Elife 12, (2023).

57. B. Tasic et al., Adult mouse cortical cell taxonomy revealed by single cell transcriptomics. Nat Neurosci 19, 335–346 (2016).

58. A. Paul et al., Transcriptional Architecture of Synaptic Communication Delineates GABAergic Neuron Identity. Cell 171, 522–539 e520 (2017).

59. H. X. Wang, W. J. Gao, Cell type-specific development of NMDA receptors in the interneurons of rat prefrontal cortex. Neuropsychopharmacology 34, 2028–2040 (2009).

60. F. Ali et al., Ketamine disinhibits dendrites and enhances calcium signals in prefrontal dendritic spines. Nat Commun 11, 72 (2020).

61. B. A. Ellenbroek, G. van Luijtelaar, M. Frenken, A. R. Cools, Sensory gating in rats: lack of correlation between auditory evoked potential gating and prepulse inhibition. Schizophr Bull 25, 777–788 (1999).

62. J. Urban-Ciecko, E. E. Fanselow, A. L. Barth, Neocortical somatostatin neurons reversibly silence excitatory transmission via GABAb receptors. Curr Biol 25, 722–731 (2015).

63. Q. Nai et al., Uncoupling the D1-N-methyl-D-aspartate (NMDA) receptor complex promotes NMDA-dependent long-term potentiation and working memory. Biol Psychiatry 67, 246–254 (2010).

64. M. S. Kruse, J. Premont, M. O. Krebs, T. M. Jay, Interaction of dopamine D1 with NMDA NR1 receptors in rat prefrontal cortex. Eur Neuropsychopharmacol 19, 296–304 (2009).

65. C. Bimbard et al., Behavioral origin of sound-evoked activity in mouse visual cortex. Nat Neurosci 26, 251–258 (2023).

66. L. Chen, C. R. Yang, Interaction of dopamine D1 and NMDA receptors mediates acute clozapine potentiation of glutamate EPSPs in rat prefrontal cortex. J Neurophysiol 87, 2324–2336 (2002).

67. H. Wang, G. G. Stradtman, 3rd, X. J. Wang, W. J. Gao, A specialized NMDA receptor function in layer 5 recurrent microcircuitry of the adult rat prefrontal cortex. Proc Natl Acad Sci U S A 105, 16791–16796 (2008).

68. H. Jaaro-Peled et al., Regulation of sensorimotor gating via Disc1/Huntingtin-mediated Bdnf transport in the cortico-striatal circuit. Mol Psychiatry 27, 1805–1815 (2022).

69. S. J. Dienel et al., Diagnostic Specificity and Association With Cognition of Molecular Alterations in Prefrontal Somatostatin Neurons in Schizophrenia. JAMA Psychiatry 80, 1235–1245 (2023).

70. R. Rajalingham et al., Chronically implantable LED arrays for behavioral optogenetics in primates. Nat Methods 18, 1112–1116 (2021).

71. A. R. Harris, F. Gilbert, Restoring vision using optogenetics without being blind to the risks. Graefes Arch Clin Exp Ophthalmol 260, 41–45 (2022).

72. T. Kuznetsova, K. Antos, E. Malinina, S. Papaioannou, P. Medini, Visual stimulation with blue wavelength light drives V1 effectively eliminating stray light contamination during two-photon calcium imaging. J Neurosci Methods 362, 109287 (2021).

73. K. B. Franklin;, G. Paxinos, The Mouse Brain in Stereotaxic Coordinates, Compact: The Coronal Plates and Diagrams (Academic Press, ed. third, 2008).

